# Growth‐mediated negative feedback shapes quantitative antibiotic response

**DOI:** 10.1101/2020.12.28.424579

**Authors:** S. Andreas Angermayr, Guillaume Chevereau, Tobias Bollenbach

## Abstract

Dose‐response relationships are a general concept for quantitatively describing biological systems across multiple scales, from the molecular to the whole‐cell level. A clinically relevant example is the bacterial growth response to antibiotics, which is routinely characterized by dose‐response curves. The shape of the dose‐response curve varies drastically between antibiotics and plays a key role for treatments, drug interactions, and resistance evolution. However, the mechanisms shaping the dose‐response curve remain largely unclear. Here, we show in *Escherichia coli* that the distinctively shallow dose‐response curve of the antibiotic trimethoprim is caused by a negative growth‐mediated feedback loop: Trimethoprim slows growth, which in turn weakens the effect of this antibiotic. At the molecular level, this feedback is caused by the upregulation of the drug target dihydrofolate reductase (FolA/DHFR). We show that this upregulation is not a specific response to trimethoprim but follows a universal trend line that depends only on growth rate, irrespective of its cause. Rewiring the feedback loop alters the dose‐response curve in a predictable manner, which we corroborate with a mathematical model. Our results indicate that growth‐mediated feedback loops shape drug responses and could be exploited to design evolutionary traps that enable selection against drug resistance.

## Introduction

Dose‐response curves are a central concept in systems biology and essential for understanding emergent nonlinear phenomena at different scales. A prime example is bacterial gene regulation where cooperativity of transcription factor binding to promoter regions governs the steepness of dose‐response curves that characterize gene expression as a function of transcription factor concentration (Bintu *et al*, 2005). The steepness of such transcription factor dose‐response curves ultimately determines whether feedback loops in genetic circuits can produce biologically relevant functions such as bistability or oscillations (Elowitz & Leibler, 2000; Gardner *et al*, 2000). At the population level, the bacterial response to antibiotics is captured by similar dose‐response curves that quantify the dependence of growth rate on drug concentration. Antibiotic dose‐response curves are routinely measured to characterize antibiotic susceptibility via the minimal inhibitory concentration (MIC) or the concentration leading to 50% growth inhibition (IC_50_), two classic quantities to describe antibiotic efficacy. However, the quantitative shape of the antibiotic dose‐ response curve – especially its steepness – and its implications are underappreciated.

The steepness of the dose‐response curve varies drastically between antibiotics. For many antibiotics, the growth rate drops gradually from high to low as the drug concentration is increased (Fig. 1A); in particular, this is the case for antibiotics targeting DNA replication at the gyrase (e.g. ciprofloxacin) or antibiotics targeting translation at the ribosome (e.g. tetracycline). Beta‐lactams like mecillinam (an antibiotic targeting cell wall biosynthesis at a penicillin binding protein) have extremely steep dose‐response curves where just a slight relative increase in drug concentration – by about two‐fold – causes an abrupt transition from full‐speed growth to near‐zero net growth (Fig. 1A). At the other end of the spectrum, the folic acid synthesis inhibitor trimethoprim (TMP) has an extremely shallow dose‐response curve (Palmer & Kishony, 2014; Rodrigues *et al*, 2016; Russ & Kishony, 2018; Chevereau *et al*, 2015): Reducing growth from full speed to zero with TMP requires a more than 100‐fold increase in drug concentration (Fig. 1A). In general, dose‐response curves are well approximated by Hill functions and the Hill slope *n* (“dose‐sensitivity”) is a quantitative measure of their steepness (Regoes *et al*, 2004; Chou & Talalay, 1983; Chevereau *et al*, 2015): TMP has *n* ≈ 1.1, while most antibiotics fall in the range 1.8 ≤ *n* ≤ 3.5, and *n* > 6 (Fig. 1A).

**Fig. 1.**
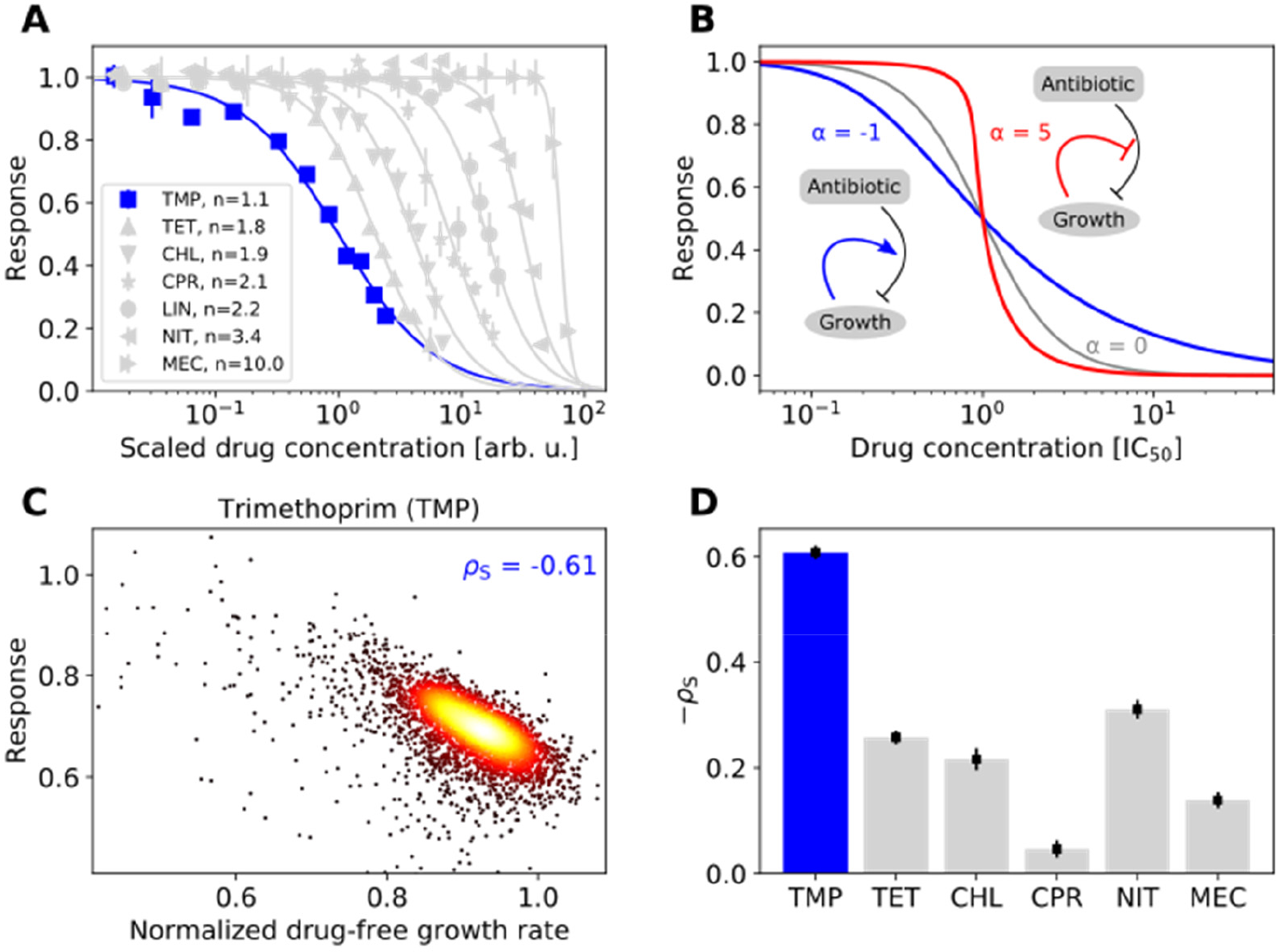
Trimethoprim exhibits an extremely shallow dose response curve and its efficacy correlates strongly with growth rate. **(A)** Dose‐response curves (normalized growth rate as a function of drug concentration) for different antibiotics. Growth rate was measured via optical density measurements over time (Methods). Antibiotics used: Trimethoprim (TMP), tetracycline (TET), chloramphenicol (CHL), ciprofloxacin (CPR), lincomycin (LIN), nitrofurantoin (NIT), and mecillinam (MEC). The TMP dose‐response curve (dark blue) is by far the shallowest. Lines are fits of the Hill function 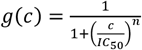 to the data. Drug concentrations were arbitrarily rescaled to better visualize dose‐response curve steepness. Error bars show standard deviation of 12 replicates. **(B)** Dose‐response curves calculated from a mathematical model that captures growth‐ mediated feedback. Negative feedback (blue) renders the dose‐response curve shallower than in the absence of feedback (gray); positive feedback (red) steepens the dose‐response curve. Parameters are n = 2, α = −1 for negative feedback and α = 5 for positive feedback (see main text); drug concentrations are normalized to the IC_50_ in the absence of growth mediated feedback (α = 0). **(C)** Density scatterplot showing growth response to TMP versus normalized drug‐free growth rate for genome‐wide gene deletion strains (Baba et al, 2006). These gene deletion strains exhibit diverse growth rates, offering an unbiased way to test the relation between the drug‐free growth rate and the response to antibiotics. Response is defined as growth rate in the presence of TMP normalized to the drug‐free growth rate of the respective deletion strain. TMP was used at a fixed concentration that inhibits wild type growth by about 30% (Chevereau et al, 2015). Spearman correlation coefficient ρ_s_ is shown. **(D)** Bar chart showing negative Spearman correlation coefficients -ρ_s_ compared across antibiotics (Supplementary Fig. 1). Error bars show bootstrap standard error of ρ_s_. TMP (blue) exhibits by far the strongest negative correlation, indicating a growth‐mediated negative feedback loop.

The steepness of the dose‐response curve strongly affects the evolution of resistance by spontaneous mutations (Hermsen *et al*, 2012; Chevereau *et al*, 2015). Resistance mutations that slightly increase the MIC provide greater fitness benefits for drugs with a steep dose‐response curve compared to drugs with a shallow curve, implying a greater chance to fix in the population. Thus, all else being equal, the rate of resistance evolution for a drug increases with the steepness of its dose‐ response curve – a trend that is observed in evolution experiments (Chevereau *et al*, 2015). This effect is strongest for drug concentrations near the IC_50_, which occur for example when populations of motile bacteria evolve resistance in spatial drug gradients (Hol *et al*, 2016; Baym *et al*, 2016). Despite their fundamental relevance for resistance evolution and bacterial responses to antibiotics, the mechanisms that shape the dose‐response curve are largely unknown.

Feedback loops mediated by growth rate may play a key role in shaping the dose‐response curve (Deris *et al*, 2013; Greulich *et al*, 2015). The action of antibiotics affects bacterial growth but the inverse is also true: Slower growing bacteria are less rapidly killed by antibiotics targeting cell wall biosynthesis (β‐lactams) (Tuomanen *et al*, 1986; Lee *et al*, 2018) and non‐growing (persister) cells are fully protected from many antibiotics (Balaban *et al*, 2004), offering a possibility to evade antibiotic treatments. Slower growth caused by nutrient limitation also affects the bacterial susceptibility to ribosome‐targeting antibiotics (Greulich *et al*, 2015). In engineered strains expressing a constitutive resistance gene, a positive feedback loop leads to high dose‐sensitivity and even bistability (i.e. co‐existence of growing and non‐growing cells) in the presence of the ribosome‐ targeting antibiotic chloramphenicol (Deris *et al*, 2013). Positive feedback occurs as faster growth leads to the upregulation of the resistance enzyme, which in turn enables even faster growth. Growth‐mediated feedback loops could more generally explain the drastic differences in dose‐ sensitivity between antibiotics (Fig. 1A) with positive feedback producing higher (Deris *et al*, 2013) and negative feedback lower dose‐sensitivity. However, such feedback loops shaping the dose‐ response curve of sensitive wild‐type bacteria have not yet been characterized.

Here, we establish that negative growth‐mediated feedback produces an extremely shallow drug dose‐response curve. Focusing on TMP, we vary bacterial growth rates by diverse environmental and genetic perturbations and show that slower growth generally lowers the susceptibility of *Escherichia coli* to this antibiotic. The molecular origin of this phenomenon lies in the expression of the drug target, which is upregulated in response to TMP but also when growth rate is lowered by other means: TMP lowers growth, which in turn reduces susceptibility to TMP. We show that synthetically reversing this feedback loop can drastically steepen the dose‐response curve. The negative feedback loop leads to a seemingly paradoxical situation where adding the antibiotic can even enhance growth under extreme nutrient limitation. Such growth‐mediated feedback loops in drug responses could be used to design evolutionary traps that invert selection for resistance.

## Results

### A minimal mathematical model shows that growth‐mediated feedback loops affect the dose‐ sensitivity of drugs

We hypothesized that a growth‐mediated negative feedback loop could explain the extreme shallowness of the dose‐response curve of TMP (Fig. 1A). As an antibiotic, TMP lowers bacterial growth (by inhibiting dihydrofolate reductase, DHFR, encoded by *folA*). If a lower growth rate in turn protects bacteria from TMP, the dose‐response curve should become shallower. To test this idea, we began by developing a mathematical model of bacterial growth in the presence of an antibiotic. We studied a minimal model in which growth‐mediated feedback is introduced by making drug efficacy dependent on growth rate. The dose‐response curve of antibiotics is captured by a Hill function 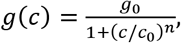, where *g*_0_ is the growth rate in the absence of the drug, *n* is the dose‐sensitivity, *c*_0_ is the drug concentration that inhibits growth by 50%, and *c* is the drug concentration, which is controlled in experiments. To capture effects of growth rate on drug efficacy, we replace *c* with an effective drug concentration *c*_eff_ that depends on the externally controlled drug concentration *c*_ext_ and on the growth rate *g*, i.e. *c* = *c*_eff_(*c*_ext_, *g*). This effective drug concentration captures that the same external concentration can have different effects on the cell, e.g. if the intracellular drug concentration or the expression level of the target is different. The resulting dose‐response curve *g*(*c*_ext_) is then implicitly given by

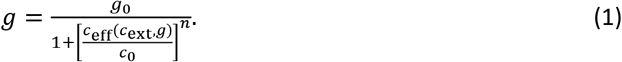

Growth‐mediated feedback is captured by a function *c*_eff_ that increases or decreases with the growth rate. For simplicity, we assume that this dependence is linear, i.e. *c*_eff_ = [1 + *α* (1 - *g*/*g*_0_)]*c*_ext_, where −1 ≤ *α* < 0 corresponds to negative and *α* > 0 to positive feedback. Solving equation (1) confirms that negative growth‐mediated feedback leads to a shallow dose‐response curve (Fig. 1B). In contrast, positive growth‐mediated feedback leads to ultrasensitivity (Fig. 1B) and can even produce bistability as previously reported (Elf *et al*, 2006; Deris *et al*, 2013). In this model, the IC_50_ also changes with *α*: IC_50_ = *c*_0_/ (1 + *α*/2). These theoretical results show how growth‐mediated feedback loops can affect the dose‐sensitivity of drugs in general.

### Slower growth generally lowers susceptibility to TMP and steepens its dose‐response curve

To test experimentally whether negative growth‐mediated feedback underlies the shallow TMP dose‐response curve, we varied growth rate in several independent ways and investigated its effect on TMP susceptibility. We first made use of a purely genetic way of varying growth. Specifically, we exploited the growth rate variability resulting from genome‐wide gene deletions to expose global trends that are independent of the specific effects of individual gene deletions. Non‐essential gene deletions often reduce the drug‐free growth rate – some by up to ~50% (Chevereau *et al*, 2015).

We re‐analyzed a dataset of growth rates of ~4,000 *E. coli* gene deletion mutants under different antibiotics representing common modes of action (Chevereau & Bollenbach, 2015); growth rates were measured at concentrations that inhibit the reference strain by ~30%. While each gene can have specific effects for each antibiotic (Nichols *et al*, 2011; Chevereau *et al*, 2015), most genes should be unrelated to the drug mode of action. The global trend of drug susceptibility across all gene deletion strains can thus reveal general consequences of growth inhibition, independent of the specific cellular limitation causing the growth rate reduction.

Non‐specific growth rate changes caused by gene deletions indicate that slower growth protects *E. coli* from TMP. By correlating the drug‐free growth rate of deletion strains with their growth rate in the presence of drugs, we revealed dependencies of drug susceptibility on the drug‐free growth rate. The clearest trend emerged for TMP: Its relative effect on growth was weaker in gene deletion strains that had lower growth rates in the absence of drugs (Spearman correlation *ρ*_s_ = −0.6; Fig. 1C). Compared to other antibiotics, this effect was most pronounced for TMP (Fig. 1D; Supplementary Fig. 1). Slower‐growing mutants can even grow at increased TMP concentrations: Full dose‐response curve measurements for 78 arbitrary gene deletion mutants showed that the IC50 is negatively correlated with the growth rate in the absence of drug for TMP (*ρ*_s_ = −0.27, *p* = 0.019) but not significantly correlated for other antibiotics (Supplementary Fig. 2). Thus, TMP represents an extreme case, both in terms of dose‐sensitivity and in terms of susceptibility‐dependence on growth rate. Overall, these results suggest that slower growth generally lowers the susceptibility to TMP.

Slow growth also protects *E. coli* from other antibiotics, but to a far lesser extent. For the prodrug nitrofurantoin (NIT) and the translation inhibitors tetracycline (TET) and chloramphenicol (CHL), there was a weak negative correlation between the drug‐free growth rate and that in the presence of the drug (*ρ*_s_ = −0.31 for NIT, *ρ*_s_ = −0.26 for TET, *ρ*_s_ = −0.21 for CHL; Fig. 1D; Supplementary Fig. 1). For the beta‐lactam mecillinam (MEC), this trend was even weaker (*ρ*_s_ = −0.14) and for ciprofloxacin (CPR) almost entirely absent (*ρ*_s_ = −0.05). Notably, the magnitude of this negative correlation roughly decreases with increasing dose‐sensitivity when compared across drugs (Fig. 1A,D). This observation supports the notion that growth‐mediated feedbacks are an important contributor to the shape of the dose‐response curve for TMP (Fig. 1B) and possibly also for other antibiotics.

Reducing growth rate by other means like nutrient limitation or imposing a protein burden also protects *E. coli* from TMP. First, we used glucose limitation in batch culture by adding a non‐metabolizable structural analog of glucose, α‐methyl glucoside, in varying concentrations to the growth medium. This analog competes with glucose for uptake into the cell, but unlike glucose it cannot be utilized for growth (Hansen *et al*, 1975). Second, we used different carbon sources (glucose, fructose, mannose, glycerol, and galactose) in the growth medium, which is a classic strategy to test for growth‐dependent effects (Bremer & Dennis, 2008). Third, we overexpressed a gratuitous protein from an inducible promoter to burden the cells (Dong *et al*, 1995; Scott *et al*, 2010). These approaches have different physiological consequences, but they all reduce growth rate in a gradual and controlled manner, while the maximal growth rate and the accessible dynamic range of relative growth inhibition varies between them (Fig. 2). Collectively, they enable us to vary growth rate over a wide range and identify general effects of growth rate, which occur independently of the exact cause of the growth rate reduction. TMP inhibits growth less under glucose limitation: Lowering growth rate by glucose limitation enabled bacteria to grow at slightly increased TMP concentrations (Fig. 2B,C). This trend was even more pronounced when growth was lowered by overexpressing a gratuitous protein – a truncated and inactive version of *tufB* (Dong *et al*, 1995) expressed from a synthetic promoter P_LlacO‐1_ (Lutz & Bujard, 1997) induced by addition of isopropyl β‐D‐1‐thiogalactopyranoside (IPTG) (Fig. 2D,E,F). Reducing growth by using different carbon sources in a minimal medium could also slightly protect bacteria from TMP, in particular for glycerol (Fig. 2G,H,I). Changing carbon sources has modest effects, presumably because even the highest growth rate (achieved with glucose only) is relatively low and the fold‐change in growth is considerably smaller than for glucose limitation (Fig. 2A,D,G). These effects did not occur to a comparable extent for other antibiotics representing common modes of action (Fig. 2C,F,I); however, gratuitous protein overexpression also lowered the susceptibility to mecillinam (MEC), albeit to a lesser extent (Fig. 2F, Supplementary Fig. 6). The effects of growth rate changes were clearly drug‐specific and strongest for TMP.

**Fig. 2.**
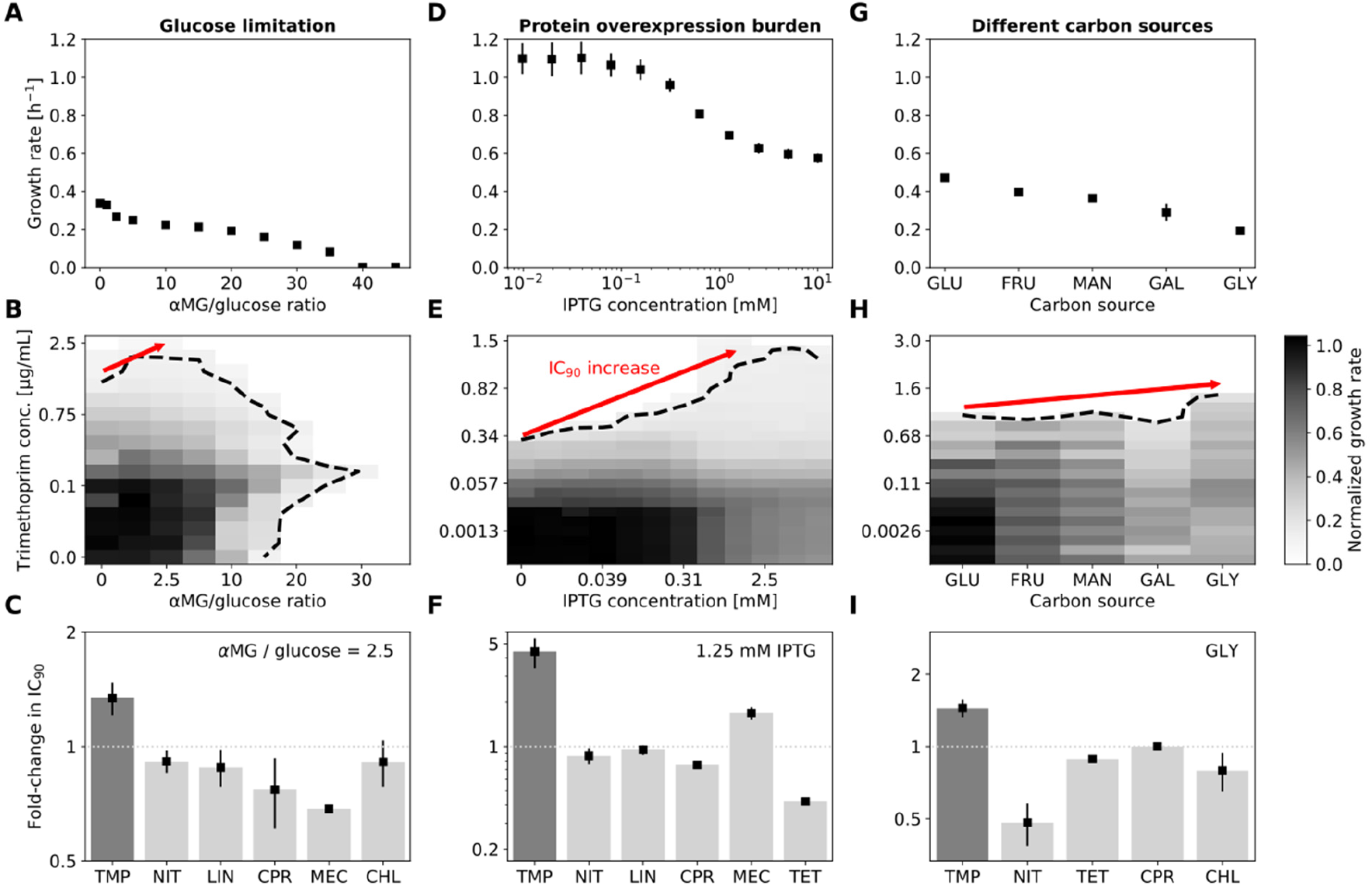
Slower growth generally lowers the efficacy of trimethoprim. **(A)** Growth rate under glucose limitation achieved by adding the non‐metabolizable structural glucose analog α‐methyl glucoside (αMG) at different ratios to glucose in a minimal medium (Methods). **(B)** Normalized growth rate (gray scale) from a checkerboard assay in a two‐dimensional concentration gradient of TMP and αMG. Dashed black line shows contour line of 90% growth inhibition (IC_90_ line). Red arrow shows increase in IC_90_ as growth is lowered. **(C)** Fold‐change in IC_90_ at αMG/glucose ratio 2.5 in assays as in B for different antibiotics (Supplementary Fig. 5). Lowering growth rate increases IC_90_ for TMP but not for other antibiotics. **(D)** Growth rate in rich medium (LB) under different levels of overexpression of a gratuitous protein from a T5‐lac promoter; overexpression burden is controlled by IPTG concentration (Methods). **(E)** As B but for growth rate reduction by protein overexpression in a two‐dimensional concentration gradient of TMP and IPTG. **(F)** Fold‐change in IC_90_ at 1.25 mM IPTG in assays as in E for different antibiotics (Supplementary Fig. 6). Overexpression of unnecessary protein increases IC_90_ for TMP by almost five‐fold; no comparable increase occurs for other antibiotics. **(G)** Growth rate in minimal medium containing different carbon sources (Methods): Glucose (GLU), fructose (FRU), mannose (MAN), galactose (GAL), and glycerol (GLY). **(H)** Normalized growth rates (gray scale) on different carbon sources (x‐axis) at different TMP concentrations (y‐axis). **(I)** Fold‐change in IC_90_ in assays as in H for different antibiotics (Supplementary Fig. 7). Error bars in A, D show standard deviation from eight replicates, those in G from three replicates; day‐to‐day reproducibility of growth rate measurements is high (Supplementary Fig. 3). Error bars in C,F show standard deviation from three neighboring αMG/glucose ratios and IPTG concentrations centered at 2.5 and 1.25 mM, respectively. IPTG alone has no detectable effect on growth at these concentrations (Supplementary Fig. 4). Error bars in I show standard deviation from three replicates. Antibiotic abbreviations are as in Fig. 1. CHL was not used in the protein overexpression assay in F since the plasmid used for overexpression has a CHL‐resistance marker (Methods). Sample growth curves are in Supplementary Fig. 11.

Under severe glucose limitation (high ratios of α‐methyl glucoside over glucose), which does not support growth, the addition of TMP even rescued bacteria and enabled them to grow again (Fig. 2B). This seemingly paradoxical phenomenon indicates that, under extreme nutrient limitation, the antibiotic TMP can facilitate bacterial growth – perhaps the most drastic illustration of the close interplay between growth rate and TMP susceptibility we observed.

Lowering growth rate by changing temperature does not show a similar effect: The relative growth reduction by antibiotics remains the same at different temperatures (Supplementary Fig. 8). This is expected since the key physiological parameters that determine growth are invariant under temperature changes (Bremer & Dennis, 2008). This observation indicates that the observed effect has a biological – rather than an elementary physical – origin.

Slower growth increases the steepness of the TMP dose‐response curve. We noticed that the extremely shallow dose‐response curve of TMP (Fig. 1A) became steeper when growth was slowed by glucose limitation (Fig. 3A): Halving the growth rate increased the dose‐sensitivity from *n* ≈ 1.1 ± 0.2 to *n* ≈ 1.6 ± 0.3 (Fig. 3B). This steepening occurred similarly when growth was slowed by gratuitous protein overexpression (Fig. 3C,D) or by changing the carbon source in the growth medium (Fig. 3E,F). In addition to this change in steepness, the concentration at which TMP starts to have an effect on growth is higher for slower‐growing bacteria. However, once the effect of TMP kicks in, the growth rate drops more rapidly with increasing TMP concentration. To better understand this unexpected increase in dose‐sensitivity resulting from slower growth, we next aimed to elucidate the underlying mechanism of TMP’s growth‐rate‐dependent action.

**Fig. 3.**
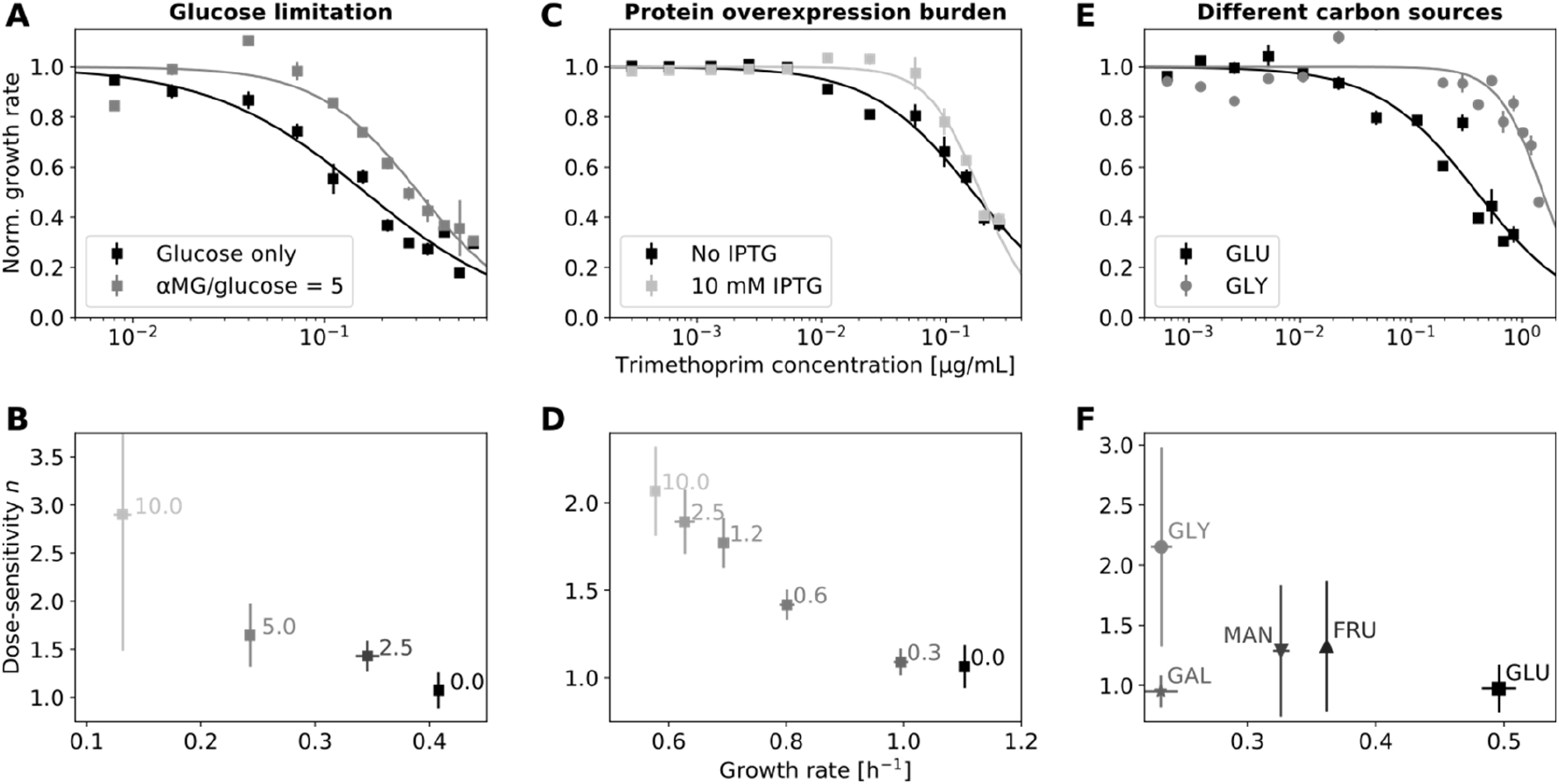
Slower growth increases the steepness of the trimethoprim dose‐response curve. **(A)** TMP dose‐response curve in minimal medium with glucose as carbon source and at lower drug‐free growth rate due to glucose limitation, achieved by increasing the αMG /glucose ratio from 0 (black) to 5 (gray). Glucose limitation results in a steeper dose‐response curve. Lines show Hill function fits (cf. Fig. 1A). **(B)** Steepness of TMP dose‐response curves (dose‐sensitivity n) versus drug‐free growth rate at different αMG concentrations (Methods). Numbers next to data points show αMG/glucose ratio. **(C,D)** As A,B but for growth limitation by gratuitous protein overexpression in rich growth medium (Methods). Inducing overexpression with IPTG at 10 mM (light gray) steepens the dose‐response curve compared to no induction (black). Numbers next to data points in D show IPTG concentration in mM. **(E,F)** As A,B but for growth limitation by varying the carbon source in a minimal medium (Methods). Carbon sources as in Fig. 2. Growth rate error bars show standard deviation of three replicates; vertical error bars in B,D,F show standard deviation of parameter estimates from Hill function fit.

### Growth‐dependent regulation of the TMP drug target leads to a negative feedback loop that flattens the dose‐response curve

Regulation of the drug target DHFR could cause the growth‐rate‐dependent efficacy of TMP. The abundance of the target of TMP (DHFR/FolA) correlates with growth (Bershtein *et al*, 2013); increasing its expression, e.g. by overexpressing *folA* from a plasmid, alleviates the effect of TMP on growth (Palmer & Kishony, 2014). Accordingly, TMP resistance in the lab and in the clinic often evolves by overexpressing *folA*, e.g. by mutating its promoter or by increasing gene copy number (Toprak *et al*, 2012; Nyerges *et al*, 2018; Rood *et al*, 1980; Flensburg & Sköld, 1987; Baym *et al*, 2016). These phenomena suggest a plausible mechanism for the reduced susceptibility to TMP at lower growth rates: We hypothesized that slower growth generally leads to increased *folA* expression, which in turn partially protects bacteria from TMP (Soo *et al*, 2011; Palmer & Kishony, 2014) – a buffering mechanism against inhibition of FolA.

DHFR expression increases similarly in response to TMP and to other means of reducing growth rate. Using a promoter‐GFP reporter (Methods), we confirmed that *folA* expression increases in response to TMP (Fig. 4A), as previously observed in whole populations (Bollenbach *et al*, 2009; Bershtein *et al*, 2015; Rodrigues *et al*, 2016) and single cells (Mitosch *et al*, 2017). However, we noticed that *folA* expression increases similarly when growth is slowed by glucose limitation (Fig. 4B). This observation suggests that the upregulation of *folA* under TMP is not a specific response to target inhibition, but rather a general response to reduced growth rate. Expression levels of constitutive genes are generally expected to increase when the quality of the nutrient environment is lowered (Scott *et al*, 2010). While the *folA* promoter can be regulated by two transcription factors (TyrR (Yang *et al*, 2007) and IHF (Keseler, 2004)) under certain conditions, it behaved similarly to a constitutive promoter in these experiments. Indeed, *folA* expression across a two‐dimensional concentration gradient of TMP and the glucose analog varied (Supplementary Fig. 9) but was largely determined by growth rate alone (Fig. 4C). Like constitutively expressed genes (Scott *et al*, 2010), *folA* expression followed a general, approximately linear increase with decreasing growth rate, approaching a fixed maximum level at zero growth (Fig. 4C). Since increased *folA* expression protects bacteria from TMP (Palmer & Kishony, 2014), this mode of regulation results in a negative growth‐mediated feedback loop: TMP inhibits growth, which leads to the upregulation of its target, thereby attenuating its own efficacy.

**Fig. 4.**
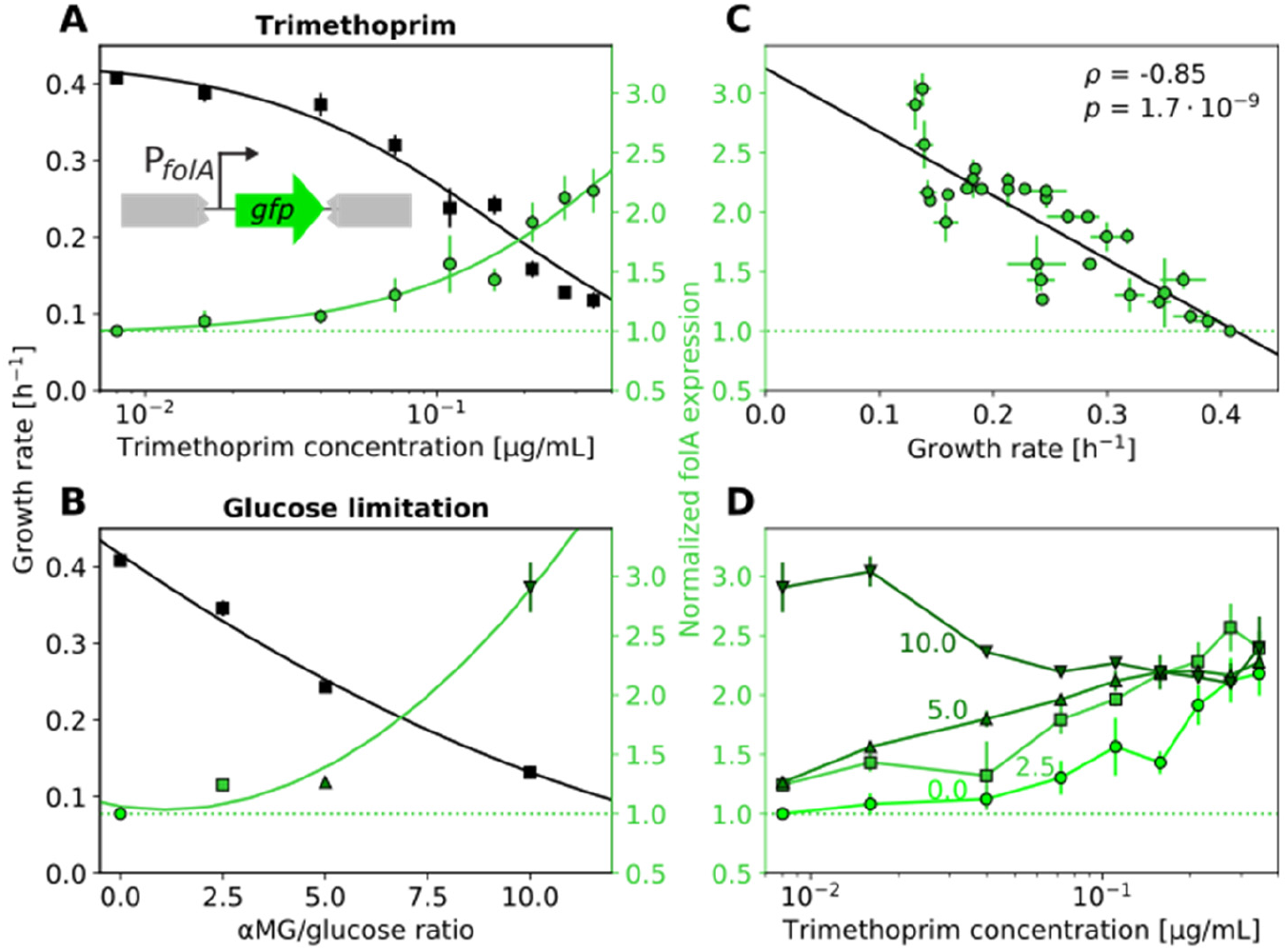
Slower growth increases folA expression, irrespective of whether growth is reduced by trimethoprim or by nutrient limitation. **(A)** Dependence of growth rate (black) and folA expression (green) on TMP concentration. Schematic: FolA expression was measured using a promoter‐GFP reporter inserted at a neutral site in the genome (Methods). **(B)** Growth rate (black) and folA expression (green) in the absence of TMP at different growth rates achieved by different ratios of αMG/glucose. **(C)** Scatterplot of folA expression level with growth rate across all combinations of TMP concentrations and αMG/glucose ratios shown in A and B (Supplementary Fig. 9). Pearson’s correlation coefficient ρ and p‐value are shown. **(D)** Dependence of folA expression on TMP concentration at four different αMG/glucose ratios (0, 2.5, 5, and 10 as shown). Darker green indicates greater αMG/glucose ratio. FolA expression converges approximately to the same level at high TMP concentrations. Black line in A shows Hill function fit as in Fig. 1A; other lines show polynomial fits of first (C) or second order (A,B) to guide the eye. Horizontal dotted green line shows folA expression level in the absence of TMP. Error bars show standard deviation of three replicates.

Saturating growth‐dependent regulation of the drug target can explain the steepening of the dose‐ response curve at lower growth rates. Higher drug target expression at lower growth rates can compensate for some of the target inhibition caused by TMP. This offers a plausible explanation as to why the effect of TMP becomes apparent only at higher concentrations when the drug‐free growth rate is lower (Fig. 2). But how does slower growth steepen the TMP dose‐response curve (Fig. 3)? We noticed that *folA* expression at different drug‐free growth rates converges to a fixed value when TMP is added (Fig. 4D). In other words, the relative upregulation of *folA* in response to TMP gets weaker with decreasing drug‐free growth rate; it even disappears completely at the highest glucose‐analog concentrations (Fig. 4D). This convergence of *folA* expression in different conditions may reflect that the promoter reaches its maximal induction level. At lower drug‐free growth rates, the promoter is already near its maximum expression level without TMP and saturates quickly when TMP is added, resulting in weaker relative upregulation than at higher growth rates. Consistent with this scenario, increasing *folA* expression at low growth rates is deleterious (Supplementary Fig. 10). Thus, lower drug‐free growth rates weaken – or even break – the growth‐ mediated negative feedback loop, resulting in steeper dose‐response curves.

### Artificially breaking the growth‐mediated feedback loop steepens the TMP dose‐response curve

To corroborate that the shallowness of the TMP dose‐response curve is due to a growth‐mediated negative feedback loop, we aimed to break this loop even under nutrient conditions that support high drug‐free growth rates. To this end, we constructed a synthetic strain in which the expression of *folA* from its endogenous locus is controlled by an inducible promoter. We used a strain allowing IPTG‐mediated induction of *folA* and *folA‐gfp,* respectively, with an expression level in the same range as wild‐type *folA* (Methods). Note that this alone does not eliminate the feedback loop since, at constant inducer levels, expression from the inducible promoter can change with growth rate, similar to expression from the endogenous *folA* promoter. Nevertheless, we can use this synthetic strain to infer the shape of the TMP dose‐response curve at constant *folA* expression by continuously varying the inducer concentration and measuring FolA levels. Specifically, we measured growth rate and *folA* expression using a FolA‐GFP fusion protein across a two‐dimensional concentration gradient of TMP and inducer (Fig. 5A; Methods). We then determined the growth rate as a function of TMP concentration on a path through this two‐dimensional concentration space along which *folA* expression is constant. The resulting TMP dose‐response curve at constant *folA* expression is steeper than in wild type (*n* = 2.0 ± 0.3; Fig. 5B,C). It becomes even steeper for a positive feedback loop, which is inferred from a path through the two‐dimensional concentration space along which *folA* expression decreases with increasing TMP concentration (*n* = 5.0 ± 0.9; Fig. 5B,C). These results provide direct evidence that a negative growth‐mediated feedback loop implemented by the regulation of the drug target causes the exceptional shallowness of the TMP dose‐response curve.

**Fig. 5.**
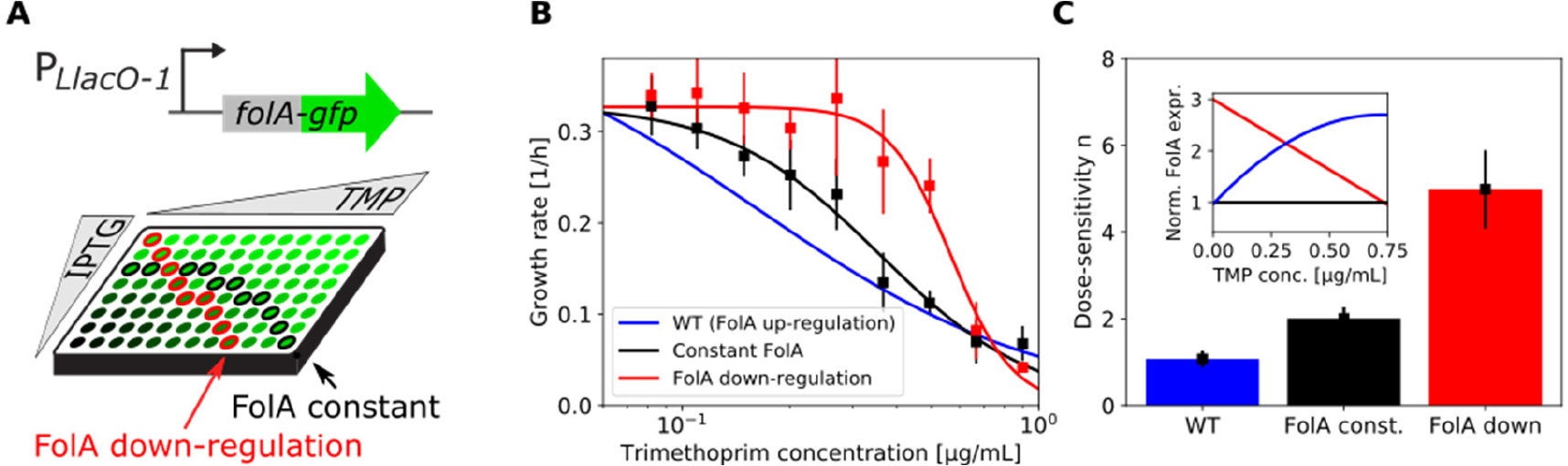
Breaking the growth‐mediated negative feedback loop steepens the trimethoprim dose‐response curve. **(A)** Schematic: FolA expression is controlled by varying the IPTG concentration and measured by flow cytometry using a GFP fusion to FolA. Shades of green indicate different FolA expression levels. Wells encircled in red indicate how the effect of FolA down‐regulation with increasing TMP concentration can be inferred; wells encircled in black illustrate the same for constant FolA expression. **(B)** Growth rate as a function of TMP concentration for different paths through IPTG‐TMP concentration space as illustrated in A (Methods). Constant FolA is shown in black and FolA down‐regulation in red. Wild type dose‐response curve (blue line; fit from Fig. 4A) is shown for comparison. **(C)** Steepness of the dose response curve (quantified as dose‐sensitivity n) for the three cases in B. Inset: Normalized FolA expression level as a function of TMP concentration for the three cases in B; WT (blue) shows fit from Fig. 4A; colors as in the bar chart and in B. Error bars in B show standard deviation of the measured growth rates used for interpolating the values shown (Methods). Error bars in C show standard deviation of parameter estimates from Hill function fit.

## Discussion

We showed that slower‐growing bacteria are generally less affected by TMP, largely regardless of what causes their slower growth (Fig. 2). This phenomenon implies a growth‐mediated negative feedback loop, which causes TMP’s extremely shallow dose‐response curve (Fig. 1A): TMP lowers growth, which in turn weakens the inhibitory effect of the drug. Mechanistically, this feedback loop is rooted in the expression level of the drug target DHFR, which is upregulated with decreasing growth rate (Fig. 4). Elimination or inversion of this feedback loop from negative to positive drastically steepens the dose‐response curve (Fig. 5). Together with recent work on ribosome‐targeting antibiotics (Deris *et al*, 2013), these results support a general role of growth‐mediated feedbacks in shaping antibiotic dose‐response curves (Fig. 1B).

Consistent with this view, the steepness of the dose‐response curve of antibiotics representing different modes of action correlates roughly with the decrease in drug susceptibility under slower growth (Fig. 1A,D). In particular, while their effect is less extreme than for TMP, the ribosome inhibitors CHL and TET also exhibit relatively low dose‐sensitivity and slightly reduced susceptibility under slower growth (Fig. 1D, Supplementary Fig. 1). The mechanism underlying this weaker growth‐mediated negative feedback for CHL and TET resembles that for TMP since their drug target, the ribosome, is upregulated in response to these drugs (Scott *et al*, 2010)—similar to DHFR in response to TMP. The prodrug NIT is an outlier: It has a relatively steep dose‐response curve (Fig. 1A) despite being less susceptible under slower growth (Fig. 1D). This may be caused by additional (unknown) mechanisms acting on top of the growth‐mediated feedback we focus on here. Identifying the molecular mechanisms underlying feedback loops or other phenomena that shape dose‐response curves will likely require detailed studies for each antibiotic or antibiotic class.

By using a mathematical model in which the effective drug concentration is growth‐rate dependent, we showed that growth‐mediated feedbacks generally affect the shape of the dose‐response curve. For TMP, the effective drug concentration is reduced due to the upregulation of DHFR: The same concentration of TMP has a weaker effect on growth at higher intracellular concentrations of DHFR. For simplicity, we assumed that the effective drug concentration changes linearly with the growth rate. This is in line with the experimental observation that DHFR is upregulated linearly with decreasing growth rate, approaching a maximum at zero growth (Fig. 4C). Decoupling the DHFR level from the growth rate by forcing it to a constant value, or even to decrease in response to TMP, results in a steeper dose‐response curve (Fig. 1B) in agreement with experimental observations (Fig. 5B,C). The model further helps to rationalize why dose‐response curves become steeper when the drug‐free growth rate is decreased, corresponding to a poorer nutrient environment. At very low drug‐free growth rates, the DHFR level becomes almost constant (Fig. 4D), effectively breaking the negative feedback loop and thus steepening the dose‐response curve, as observed experimentally (Fig. 3). Note, that our model can explain relative changes in dose‐sensitivity but it remains challenging to explain its absolute value. Indeed, all antibiotics we studied have dose‐sensitivities *n* > 1 and most have *n* > 2 (Fig. 1A), which requires additional nonlinearities beyond the growth‐mediated one we focused on here, e.g., in the kinetics of antibiotic transport and target binding.

Explaining the absolute values of the dose‐sensitivities for different drugs remains a challenge for future work.

We observed that an artificial nutrient limitation that results in no or extremely slow growth can be alleviated by adding the antibiotic TMP (Fig. 2B). This indicates that bacteria may not regulate DHFR expression in a way that maximizes growth under extreme nutrient limitation. Indeed, slow growth in these conditions coincides with high *folA* expression (Fig. 4D). High *folA* expression is deleterious (Bhattacharyya *et al*, 2016) as cellular resources are diverted toward excessive folic acid synthesis. Consistent with this view, TMP facilitated bacterial growth when *folA* was overexpressed to levels that were deleterious in the absence of TMP (Supplementary Fig. 10). Together, these observations show that DHFR level is the main driver of TMP susceptibility and suggest that costly overproduction of DHFR, which can be rescued by adding TMP, occurs under extreme nutrient limitation. Since TMP increases the fitness of bacteria that evolve under extreme nutrient limitation, the usual selection pressure for antibiotic resistance is inverted under such conditions: Mutations that usually enhance TMP action (e.g. increased drug uptake) can be selected. Similar to certain drug combinations (Chait *et al*, 2007), this situation provides an opportunity to select against antibiotic‐resistant bacteria. One potential advantage of creating such conditions with a sugar analog instead of a second drug is that bacteria can hardly evolve resistance to such an analog, as impaired sugar uptake would come at a massive fitness cost.

## Methods

### Growth conditions and growth rate measurements

Unless otherwise noted the chemicals used were from Sigma‐Aldrich (Steinheim, DE). The growth medium used was either LB Broth Lennox (L3022), pH set to 7.0 with NaOH before autoclaving, or M9 minimal medium made from Na_2_HPO_4_.7H_2_O (Fisher Scientific Acros Organics 206515000), KH_2_PO_4_ (P9791), NaCl (S3014), and NH_4_Cl (A9434)) supplemented with 0.1 mM CaCl2 (Fluka 223506), 2mM MgSO4 (M7506), and 0.001% (v/v) Triton‐X 100 (T8787). Triton‐X was added to flatten the meniscus that forms in 96‐well pates (Mitosch *et al*, 2017). Carbon sources in the M9 medium were glucose (G8270), glycerol (VWR 0854), mannose (Carl Roth 4220.2), fructose (F0127), galactose (G0750), all of which were added at 0.4% (w/v) and prepared as filter sterilized 20% (w/v) stock solutions stored at room temperature in the dark. Experiments were started from a frozen glycerol stock. Bacteria were streaked on an LB agar (L2897) plate (containing antibiotics as appropriate) incubated overnight at 37°C and a single colony was inoculated in 2 ml of the appropriate growth medium (containing antibiotics if appropriate) and grown for about 20 h to obtain a pre‐culture that has reached stationary phase. We inoculated experimental cultures with a 1000‐fold dilution from a stationary phase culture when growth was determined by optical density measurements at 600 nm (OD_600_). For the experiments with luminescence‐readout, the pre‐culture was grown in 20 ml LB medium in a 250 ml flask until stationary phase; 100 μl aliquots were transferred to the wells of a 96‐well plate, supplemented with glycerol to 15% and frozen at ‐80°C. To start a luminescence‐based experiment the plate was thawed, and dilutions were performed in 96‐well plates with fresh medium using pin tools (VP407 and VP408, V&P Scientific Inc., CA, USA), which transfer 1.5 μl and 0.2 μl per well, respectively. Subsequent use resulted in a 10^7^‐fold dilution from a stationary phase culture. In all cases, the pre‐cultures were incubated at 30°C with a shaking speed of 250 rpm (Innova 44, Eppendorf New Brunswick, DE).

Pre‐cultures carrying plasmids and cultures needed for molecular cloning procedures were prepared with antibiotics at the following concentrations: chloramphenicol 35 μg/ml (C0378), kanamycin 25 μg/ml (K4000), ampicillin 50 μg/ml (A9518), spectinomycin 100 μg/ml (S6501).

Unless otherwise noted antibiotics were dissolved in ethanol (32221). Stock solutions in water were filter‐sterilized. Aliquots of stocks were stored at ‐20°C in the dark. The antibiotics used were trimethoprim (92131), nitrofurantoin (N7878), chloramphenicol (C0378), lincomycin (dissolved in water, 62143), mecillinam (dissolved in water, 33447), tetracycline (268054), and ciprofloxacin (dissolved in water, 17850). IPTG (VWR 437144N) was added to cultures to control expression from IPTG‐responsive promoters (P_T5‐lac_, P_LlacO‐1_) (Kitagawa *et al*, 2005; Lutz & Bujard, 1997). A filter‐sterilized solution of 1M IPTG in water served as stock solution. IPTG was stored at ‐20°C in the dark and aliquots were thawed at room temperature before use. For the non‐metabolizable glucose analog α‐methyl glucoside (M9376), which competes for glucose uptake and essentially imposes glucose limitation (Hansen *et al*, 1975), a filter sterilized solution of 50% (w/v) in M9 salts served as stock solution.

The experiments shown in Fig. 1‐3 were performed using a robotic system as described previously (Chevereau *et al*, 2015) and have a day‐to‐day variability (coefficient of variation, CV) of growth rate for unperturbed cultures of less than 5% (Ref. (Chevereau *et al*, 2015) and Supplementary Fig. 3). The experiments shown in Fig. 4 and 5 were performed using two plate readers: A Synergy Neo2 and a Synergy H1 (both from Biotek Inc., VT, USA). Both were set to 30°C with continuous shaking at an orbital displacement of 1 mm and a speed of 807 rpm, and after a settling period of 10 seconds the optical density at 600 nm and GFP fluorescence were measured every 10 min. Flat transparent microtiter plates (Nunc Thermo Scientific FT 96‐well, 236105) with lids were used. The experiments presented in Supplementary Fig. 8 were performed using an Infinite M1000 Pro plate reader (Tecan Inc., CH) equipped with an integrated stacking module. The stack was housed in a custom‐built (IST Austria Miba Machine Shop, Klosterneuburg, AT) acrylic glass box equipped with a custom‐built heating block, a thermostat and strong ventilation to assure a homogenous temperature over the plates and the stack (Kavčič *et al*, 2020). For these experiments (Supplementary Fig. 8), the wild‐type strain used here (*E. coli* BW25113) was transformed with a kanamycin resistance‐bearing plasmid (pCS‐λ) carrying luciferase genes used to determine the growth rate (Kishony & Leibler, 2003; Chait *et al*, 2007). For the actual growth experiments kanamycin was omitted; however this was not a problem as the plasmid is retained throughout the duration of such an experiment (Kavčič *et al*, 2020). Luminescence assays were performed using flat white microtiter plates (Nunc Thermo Scientific FW 96‐well, 260860). These plates were sealed with a transparent foil (TopSeal‐A Plus, PerkinElmer) and about ten plates were used per stack. Luminescence was measured every 10 to 20 min. Before each measurement, plates were shaken for 10 sec at 582 rpm with a 1 mm amplitude. The culture volume per well was 150 μl. The day‐to‐day CV for unperturbed cultures for the growth rate in the luminescence‐setup was 3%.

The growth rate was determined by a linear fit of the log‐transformed and background‐subtracted OD_600_ from the exponential growth phase of the cultures using custom Matlab (R2016b, MathWorks Inc.) scripts. To capture the exponential growth phase for cultures in LB we used background‐subtracted OD_600_ windows of 0.02 to 0.2 and for minimal medium 0.03 to 0.12; these windows cover one order of magnitude and at least two doublings and take the lower growth yield in minimal medium into account. The lowest accepted growth rate for LB was 0.1 h^‐1^ and for minimal medium 0.03 h^‐1^, both corresponding to about 10% of the respective unperturbed maximal growth rate. For the Hill function fits in Fig. 3C,D, growth rates below 0.2 h^‐1^ were ignored because too many data points fell in this range at higher IPTG concentrations (Fig. 2E) – these data points would thus dominate the fit, which is undesirable since they contain less information about the shape of the dose‐response curve (i.e. the dose‐sensitivity). The duration of experiments for LB cultures was about 22 h, for minimal medium about 46 h. For all experiments performed in LB medium, data after ~1,000 min were discarded to avoid the inclusion of faster growing mutants which occurred sporadically in the presence of antibiotics; this was not necessary for experiments in minimal medium. To capture the growth rate strictly during exponential phase from the luminescence‐based experiments, the rate of luminescence increase was determined by a linear fit of the log‐transformed data between 10^2^ cps and 10^5^ cps.

### Expression level measurements using plate readers

Two plate readers Synergy Neo2 and a Synergy H1 (see section *Growth conditions* for further details) were used for GFP fluorescence measurements. The filter set used in the Neo2 provided excitation at 485nm (BW20) and emission at 516nm (BW20) (Biotek fluorescent filter #105). The settings for the monochromator‐based H1 model were 485 nm for excitation and 528 nm for emission. Both readers produced consistent values and results. The measured values for experiments where both plate readers were used in parallel were adjusted accordingly (i.e. simply normalized by a constant obtained from measuring the same sample on both readers). The expression level was determined essentially as described (Zaslaver *et al*, 2006; Mitosch *et al*, 2017). Briefly, for each GFP‐expressing strain, a similar strain without GFP‐expression was grown in parallel in the same conditions (see section *Strain construction* for further details). For both strains the exponential growth phase was determined and the background subtracted GFP‐signal from the GFP‐less strain was subtracted from the GFP‐carrying strain for cultures with similar growth rates and at the same OD_600_. As the exact same OD_600_ values were mostly not met, linear interpolation (Matlab function *interp1*) was used to generate an interpolated GFP‐value between the two GFP values of the two nearest OD_600_ values. The expression level is obtained from the slope of a linear fit (Matlab function *fit*) to the GFP over OD_600_ data during exponential growth. In the experiments using the strains with the reporter construct with the native promoter (BWAA01, Fig. 4), fast‐folding GFP (Zaslaver *et al*, 2006) was used whereas in the experiments with the synthetic IPTG‐inducible promoter construct (BWAA29R1, Fig. 5) the GFP from the ASKA‐library (Kitagawa *et al*, 2005) was used.

### Expression level measurements using flow cytometry

For the expression level determination of the strains with IPTG‐induced *folA‐gfp* expression (Fig. 5) we used a combination of plate readers (Biotek Synergy H1) for optical density measurements for growth rate determination (see section growth rate measurements for details) and flow cytometry (Beckman Coulter CytoFLEX B2‐RQ‐V2 with 96‐well plate module) for fluorescence measurements. Flow cytometry was used because of its higher signal‐to‐noise ratio compared to fluorescence measurements on plate readers. Strains were grown in the plate readers and growth was monitored by measuring optical density every 10 min. When strains were in mid‐exponential growth phase (OD ~ 0.1), they were diluted 1,000‐fold in ISOTON II (Beckman Coulter) and measured immediately on the flow cytometer. Gating in SSC‐A and GFP FITC‐A channels in the flow cytometry analysis software (Beckman Coulter Cytexpert 2.3.0.84) allows finding of (fluorescent) cells and determination of the mean and relative coefficient of variation of fluorescence intensity. Strains used were BWAA11, BWAA 12, BWAA 19R1, and BWAA 20R1 (see section *Strain construction* for details) and TMP and IPTG gradients starting at 0.9 μg/ml and 2.5 mM were applied, respectively. Growth rates at constant or decreasing FolA expression level were calculated by linear interpolation of the growth rates measured at different IPTG and TMP concentrations as illustrated in Fig. 5A.

### Strains and strain construction

We used *E. coli* BW25113 and several derivatives thereof. BW25113 is the parent strain of the KEIO collection, a widely used whole‐genome deletion mutant collection (Baba *et al*, 2006). For the overexpression experiments, BW25113 was transformed with the necessary plasmids (Table 1) which stem from the ASKA‐library, a plasmid‐based whole‐genome overexpression collection (Kitagawa *et al*, 2005). To reduce growth rate by gratuitous protein expression we used a truncated elongation factor Tu (EF‐Tu, *tufB*) as previously done for a similar purpose (Dong *et al*, 1995). Briefly, starting with the ASKA‐library plasmid carrying *tufB*, the SmaI restriction fragment of 243 bp in length was cut out and the blunt‐ended DNA fragment was closed by ligation to form a plasmid again, named pAAtufB here. This deletion results in a shortened, non‐functional gene (*ΔtufB*), which can be used to provide gratuitous protein expression, resulting in a burden that slows down growth (Dong *et al*, 1995; Scott *et al*, 2010). The plasmids from the ASKA‐library (Kitagawa *et al*, 2005) use the P_T5‐lac_ promoter, which allows for a graded control of expression by the addition of the inducer IPTG (which works sufficiently well in a *lac*‐operon compromised strain like *E. coli* BW25113). As control, we used pAA30 which is the empty ASKA plasmid modified to not contain a gene to prevent any expression; we created this plasmid since the original empty ASKA plasmid does in fact encode a short coding sequence in frame with the promoter. Briefly, through a PCR with overlapping primers (for general strategy see (Heckman & Pease, 2007; Hansson *et al*, 2008)) (Fw: CATTAAAGAGGAGAAATTAACTGGGTCGACCTGCAG, Rv: CTGCAGGTCGACCCAGTTAATTTCTCCTCTTTAATG) a short stretch of pCA24N(‐) encompassing start codon over the His‐Tag and until the stop codon, was eliminated. The elimination was confirmed by sequencing the resulting plasmid with primers flanking the gene insertion site (Fw: CAACAGTTGCCTAAGAAACCAT, Rv: TGAGGTCATTACTGGATCTATCAAC). For the strong *folA* overexpression the ASKA plasmid pCA24N(‐)folA was used.

We generated reporter strains and a strain with inducible *folA* regulation. To construct the first *gfp*‐ reporter and corresponding *gfp*‐less control pair integrated into the chromosome (BWAA01 and BWAA02), the promoter‐reporter construct for P_folA_ and the corresponding region from the empty plasmid pUA66 from the reporter library (Zaslaver *et al*, 2006) were integrated into a neutral site (*phoA*) in the genome, respectively. To this end, P1 transduction was used to move the construct from an MG1655 strain carrying the reporter constructs (Bollenbach *et al*, 2009) into the BW25113 background. The insertion was confirmed by sequencing PCR products generated using primers binding outside the *phoA* locus (Fw: GGCGCTGTACGAGGTAAAG, Rv: GGGTTAAAGTTCTCTCGGCA).

The other reporters were based on *folA‐gfp* fusion constructs from the ASKA‐library (Kitagawa *et al*, 2005). Again, pairs of strains were made where each pair consists of a strain with and a strain without the *gfp* fused to *folA*. We generated several pairs to induce and thereby control expression level by an IPTG‐responsive promoter (P_LlacO‐1_ (Lutz & Bujard, 1997)) and one pair with the native regulation through P_folA_. This was done to approximately match the induced expression with the level of the native regulation. We generated strains with the inducible P_LlacO‐1_ promoter with five different ribosome binding sites of different strength with the sequences (RBS1, 3, 4, 5, and 6) originating from (Deris *et al*, 2013). Those strains were compared to strains with the native regulation (BWAA11 and BWAA12) which were made in parallel according to a similar strategy as described below. Based on similar expression levels for BWAA19R1 and BWAA11, BWAA19R1 with RBS1, was chosen for the experiment shown in Fig. 5. To create the *gfp* fusion strains, *folA‐chlR* and *folA‐gfp‐chlR* fragments were PCR‐amplified from the *folA*‐carrying ASKA‐library plasmids (using as template the respective plasmids from the library, with and without *gfp* (Kitagawa *et al*, 2005)) as a first step and were used for recombineering (Datsenko & Wanner, 2000) into the plasmid pKD13‐gfpmut3 (a derivative of pKD13 (Datsenko & Wanner, 2000); gift from Bor Kavčič). Primers used were CAGCAGGACGCACTGACCGAATTCATTAAAGAGGAGAAAGGTACCGCATGATCAGTCTGATTGCGGCGTTAG and GACTGAGCCTTTCGTTTTATTTGATGCCTCTAGACTCAGCTAATTAAGCGTAGCACCAGGCGTTTAAGG. This resulted in pAA39 and pAA40 where an FRT‐flanked kanamycin resistance cassette, the promoter P_*LlacO‐1*_ driving *folA* and the *folA‐gfp* fusion, respectively, and a chloramphenicol resistance cassette are present (in this order). These plasmids first served as source for the promoter‐*folA* and *folA‐gfp* fusion with a chloramphenicol resistance cassette to be inserted into the genome at the *folA* locus to generate the strains with the native regulation (BWAA11 and BWAA12). PCR‐fragments for recombineering were obtained with the previously used forward primer CAGCAGGACGCACTGACCGAATTCATTAAAGAGGAGAAAGGTACCGCATGATCAGTCTGATTGCGGCGTTAG and AAGACGCGACCGGCGTCGCATCCGGCGCTAGCCGTAAATTCTATACAAAACTAGACTCAGCTAATTAAGC serving as reverse primer to get the *folA* gene with and without *gfp,* respectively, and the chloramphenicol resistance cassette (but not the synthetic promoter) were inserted into the genome of BW25113 replacing the *folA* gene (but not the promoter on the genome). Next, by recombineering with PCR‐fragments containing the kanamycin resistance cassette only obtained with the primers GTGATACGCCTATTTTTATAGGTTAATGTCATGATAATAATGGTTTCTTAGAAGTTCCTATTCTCTAGAAAGTAT AGG and AAGACGCGACCGGCGTCGCATCCGGCGCTAGCCGTAAATTCTATACAAAAGTGTAGGCTGGAGCTGCTTC from pKD13‐gfpmut3 the chloramphenicol resistance cassette was replaced with the FRT‐flanked kanamycin resistance cassette. Next, to obtain a marker‐less strain the kanamycin resistance cassette was removed using the plasmid pCP20, as described (Cherepanov & Wackernagel, 1995). For the strains with the IPTG‐inducible regulation (BWAA19R1 and BWAA20R1) a similar strategy applied. Primers AACATACCGGCAACATGGCGGATGAACCGGAAACGAAACCCTCATCCTAAGTGTAGGCTGGAGCTGC and TCCATGCCGATAACGCGATCTACCGCTAACGCCGCAATCAGACTGATCATGCGGTACCTTT<u>CT</u>CCTCTTT (putative RBS1 sequence from (Baba *et al*, 2006) underlined) were used to amplify the FRT‐flanked kanamycin resistance cassette, the promoter P_*LlacO‐1*_ driving *folA* and the *folA‐gfp* respectively of pAA39 and pAA40 (but not the chloramphenicol resistance cassette). Next, to obtain a marker‐less strain the kanamycin resistance cassette was removed using pCP20. All four marker‐less strains were further modified by P1 transduction from a MG1655 strain carrying the *lacI* gene under the promoter P_*lacO1*_ and a FRT‐flanked kanamycin resistance cassette at the neutral insertion site *intS* (based on the strain from (Garcia *et al*, 2011) (HG105) and a gift from Bor Kavčič). The insertion was confirmed by sequencing PCR products generated using primers binding outside the *intS* locus (Fw: GTACTTACCCCGCACTCCAT, Rv: TGTTCAGCACACCAATAGAGG). Next, that kanamycin resistance cassette was removed using pCP20. The resulting markerless strains were further modified by P1 transduction with the *lacI* knock‐out strain from the KEIO collection (Baba *et al*, 2006) replacing the *lacI* gene with the kanamycin resistance cassette. The deletion was confirmed by sequencing PCR products generated using primers binding outside the *lacI* locus (Fw: CGGCTCATGGATGGTGTT, Rv: CGAAGCGGCATGCATTTAC). We reasoned, that here a P_L*lacO‐1*_‐driven *lacI* allows a better control of the P_L*lacO‐1*_‐driven *folA* based on observations in (Kavčič *et al*, 2020; Klumpp *et al*, 2009) dealing with growth rate independent negative autoregulation. Moreover, with the combination of RBS1 and the Lac‐repressor driven by P_*lacO1*_ we achieved expression close to wild‐type levels. For the generation of the strains with modified RBS3‐6 from BWAA19R1 and BWAA20R1, the kanamycin resistance cassette was removed using pCP20. We again used long reverse primers RBS3‐Rv: TCCATGCCGATAACGCGATCTACCGCTAACGCCGCAATCAGACTGATCATGCGGTACCTTT<u>AGGACT</u>CCTCTTT aatgaattcggtcag, RBS4‐Rv: TCCATGCCGATAACGCGATCTACCGCTAACGCCGCAATCAGACTGATCATGCGGTACCTTT<u>GT</u>CCTCTTTaatga attcggtcag, RBS5‐Rv: TCCATGCCGATAACGCGATCTACCGCTAACGCCGCAATCAGACTGATCATGCGGTACCTTT<u>AGGAGT</u>CCTCTTT aatgaattcggtcag, RBS6‐Rv: TCCATGCCGATAACGCGATCTACCGCTAACGCCGCAATCAGACTGATCATGCGGTACCTTT<u>AGGAGG</u>CCTCTTT aatgaattcggtcag, each modified such that the RBS sequence was changed according to (Deris *et al*, 2013). To aid primer binding (compare (Liu & Naismith, 2008)) to the template (which carries RBS1 sequence) and change of the RBS‐sequence (underlined) additional 15 nt were added to the reverse primers (lower case letters) and used in combination with Fw: AACATACCGGCAACATGGCGGATGAACCGGAAACGAAACCCTCATCCTAAGTGTAGGCTGGAGCTGC on pAA39 and pAA40. The respective PCR products were sequenced and used for recombineering of the markerless intermediate versions of BWAA19R1 and BWAA20R1. After integration, all promoter regions were confirmed by sequencing a PCR product obtained with primers targeting the *folA* locus with the primers (Fw: CCAGCGCGATGTTAAGTGA, Rv: GATTGATTCCCAGGTATGGCG).

For the recombineering procedure (Datsenko & Wanner, 2000) the temperature‐inducible system from pSIM19 (Sharan *et al*, 2009) was used. Chloramphenicol at 10 μg/ml and kanamycin at 25 μg/ml were used. During the whole strain construction procedure wherever *folA* was driven by PL_*lacO1*_, 1 mM IPTG was added as this inducer controls expression from PL_*lacO1*_ and *folA* is an essential gene.

### Mathematical model

Solutions of the mathematical model in Fig. 1B were numerically calculated using Python (function *fsolve* from *scipy* version 1.2.1). As dose‐sensitivity without growth‐mediated feedback, we used *n* = 2 since this is a typical value for many antibiotics (Fig. 1A).

### Strains and plasmids used in this study

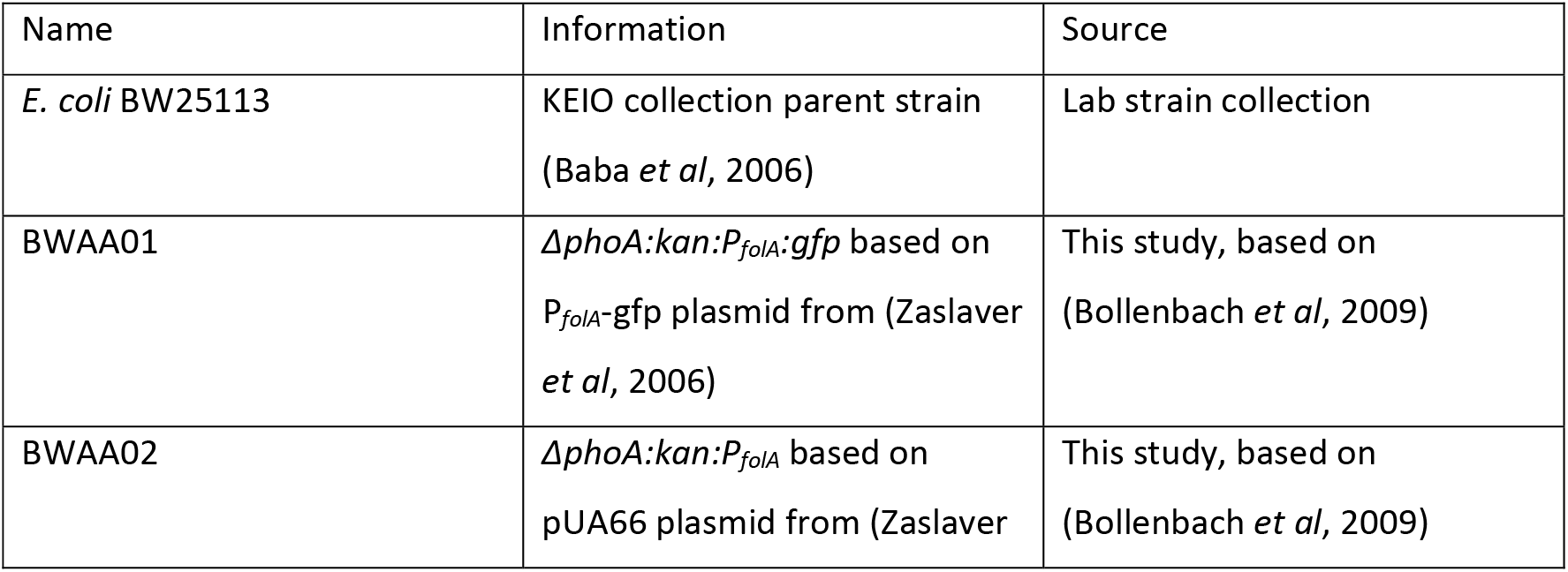
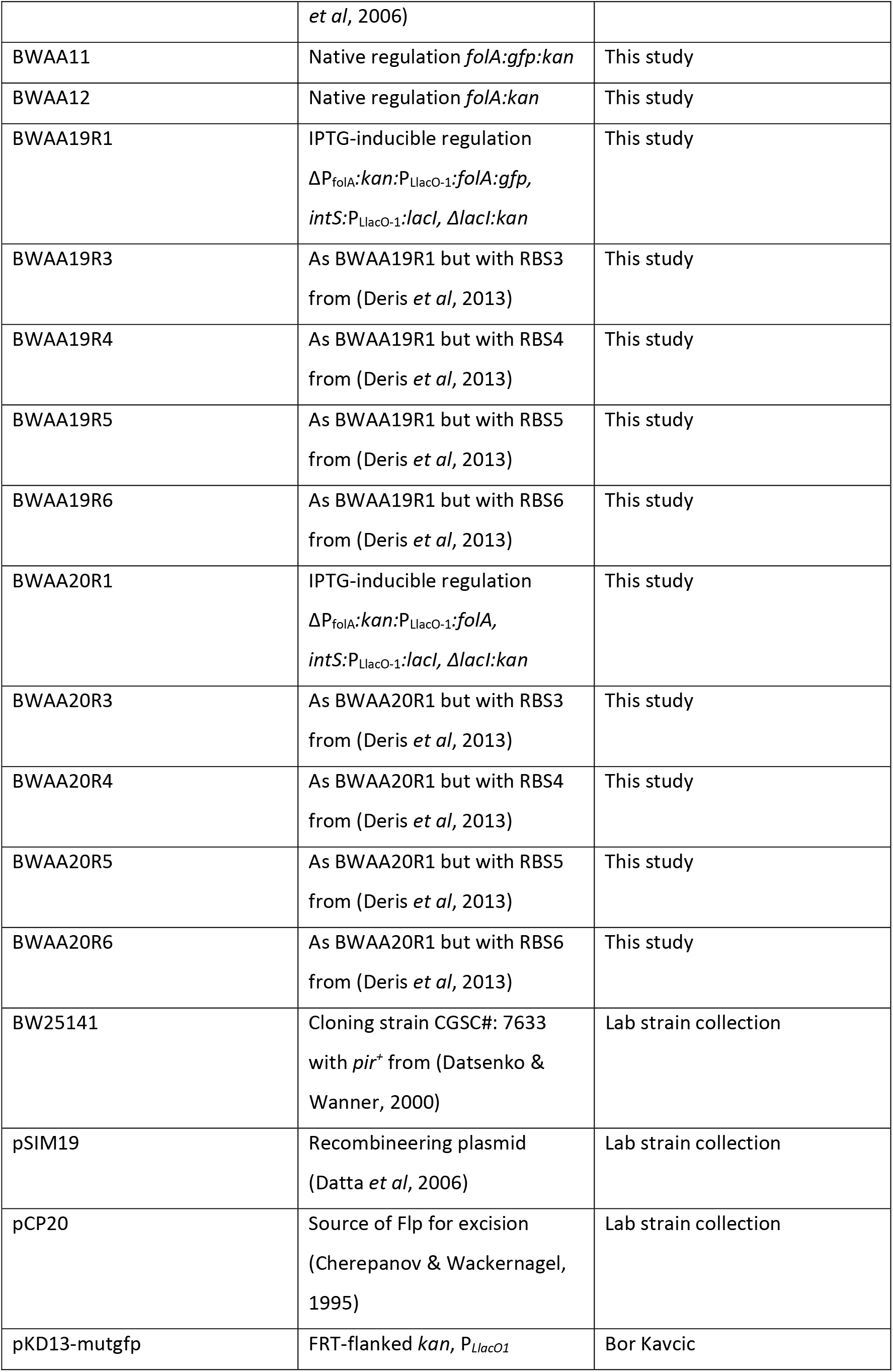
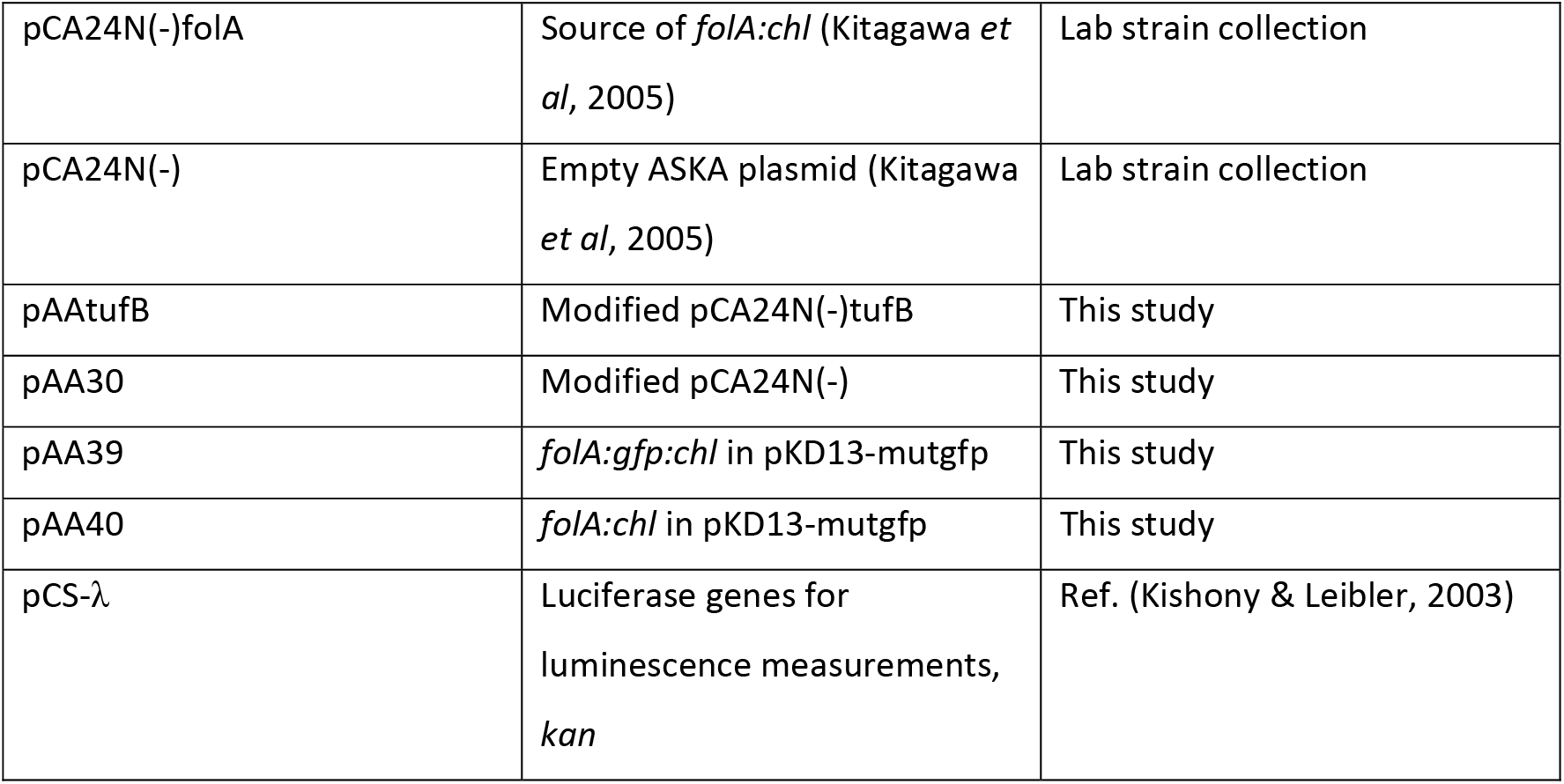

## Acknowledgements

This work was in part supported by Human Frontier Science Program Grant RGP0042/2013, Marie Curie Career Integration Grant 303507, Austrian Science Fund (FWF) Grant P 27201‐B22, and German Research Foundation (DFG) Collaborative Research Center (SFB) 1310 to Tobias Bollenbach. S. Andreas Angermayr was supported by the European Union’s Horizon 2020 research and innovation program under the Marie Skłodowska‐Curie grant agreement No 707352. We would like to thank the Bollenbach group for regular fruitful discussions. We are particularly thankful for technical assistance of Booshini Fernando and for discussions of the theoretical aspects with Gerrit Ansmann. We are indebted to Bor Kavčič for invaluable advice, help with setting up the luciferase‐ based growth monitoring system, and for sharing plasmids. We are grateful to Rosalind Allen, Bor Kavčič, Karin Mitosch and Dor Russ for feedback on the manuscript.

We declare that there are no conflicts of interest.

SAA, GC, and TB conceived the study. SAA and TB designed the experiments. SAA performed the experiments and analyzed the data. GC analyzed data presented in Fig. 1C and Supplementary Fig. 1. SAA and TB wrote the manuscript with input from GC.

## Supplementary figures

**Supplementary Fig. 1.**
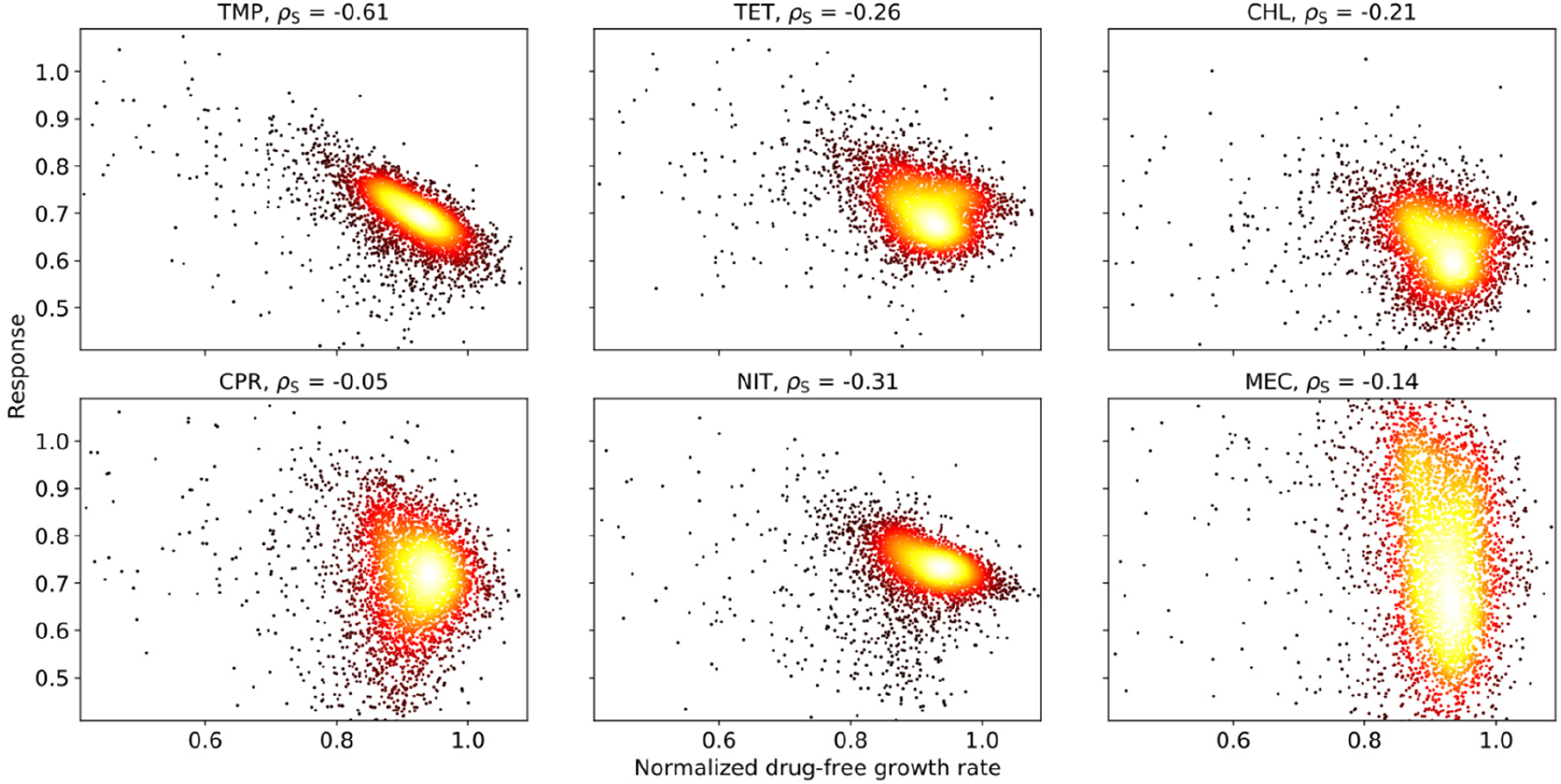
Correlation of antibiotic response and drug-free growth rate for genome-wide gene deletion strains. Density scatterplots showing growth response to different antibiotics versus normalized growth rate in the absence of drug for genome-wide gene deletion strains (Baba et al, 2006) as in Fig. 1C. Response is defined as growth rate in the presence of the respective drug normalized to the drug-free growth rate of the respective deletion strain. Each drug was used at a fixed drug concentration that inhibits wild type growth by about 30% (Chevereau et al, 2015). Spearman correlation coefficient ρ_s_ is shown.

**Supplementary Fig. 2.**
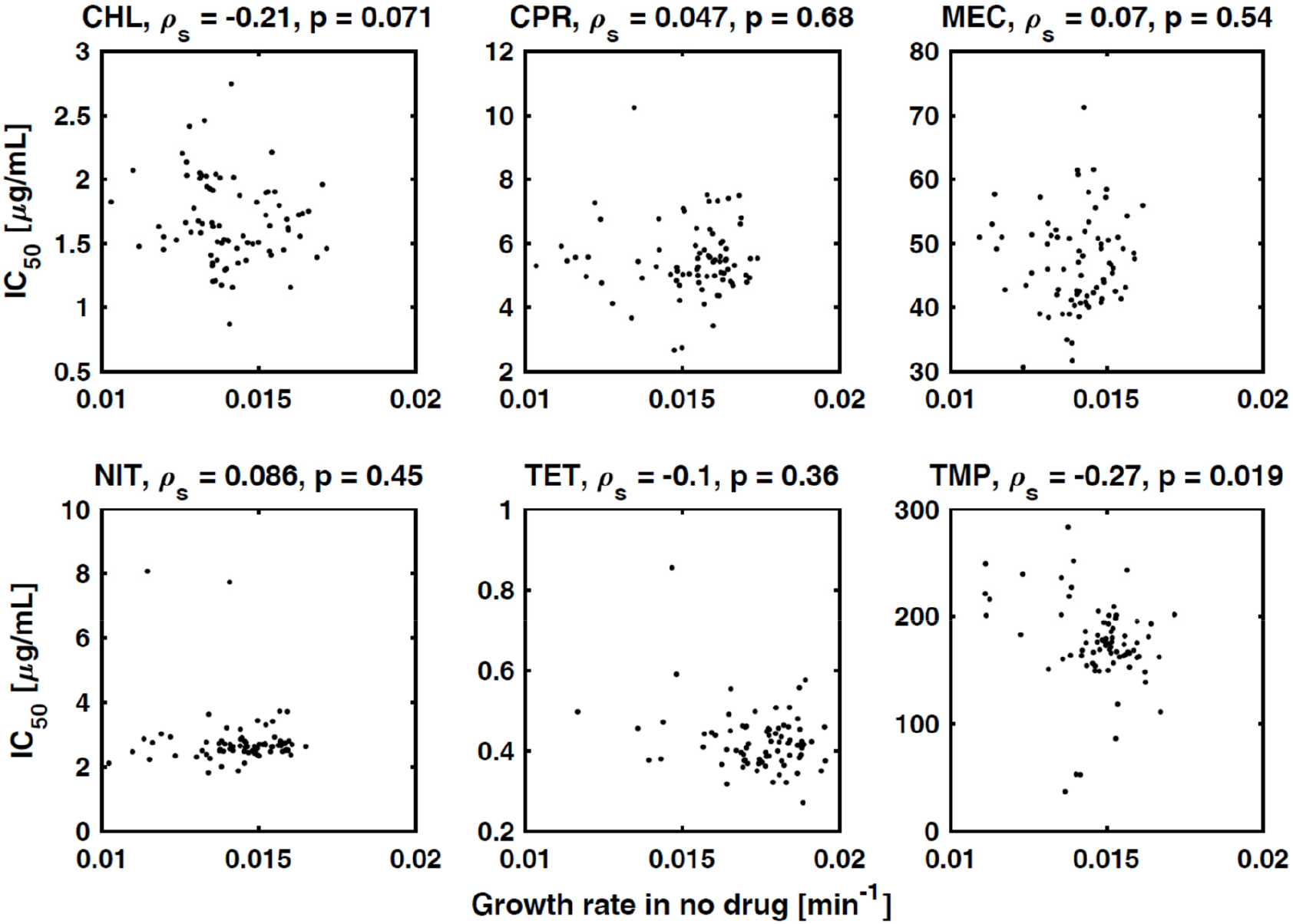
Correlation of IC_50_ and drug-free growth rate in gene deletion strains. Scatterplots of the IC_50_ of gene deletion mutants versus the growth rate of these mutants in the absence of drug; each panel shows a different antibiotic as labeled. ρ_s_ is the Spearman correlation; p-values of this correlation are shown. The only significant (negative) correlation occurs for TMP, consistent with growth-mediated negative feedback for this drug. IC_50_s were determined from dose-response curve measurements of 78 arbitrary gene deletions strains (Chevereau et al, 2015).

**Supplementary Fig. 3.**
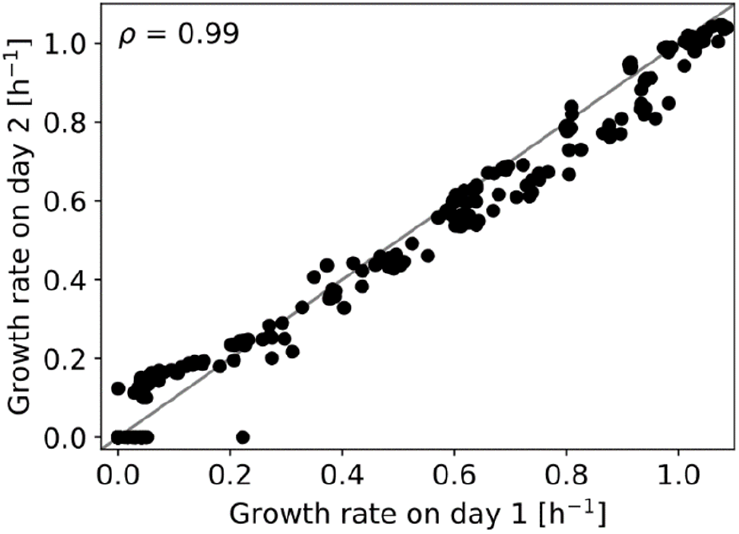
Day-to-day reproducibility of growth rate measurements. Scatterplot showing comparison of growth rate data from αMG-TMP two-dimensional concentration gradient experiment (Fig. 2B) performed on two different days.

**Supplementary Fig. 4.**
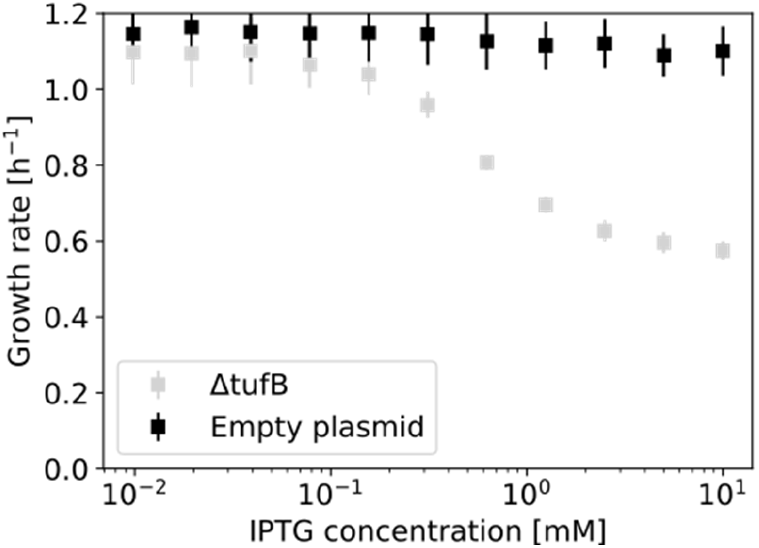
IPTG alone at the concentrations used here has no effect on growth rate. Black data points show growth rate versus IPTG concentration for a control strain with an empty expression vector; data from Fig. 2C is shown in gray for comparison. Error bars show standard deviation from eight replicates.

**Supplementary Fig. 5.**
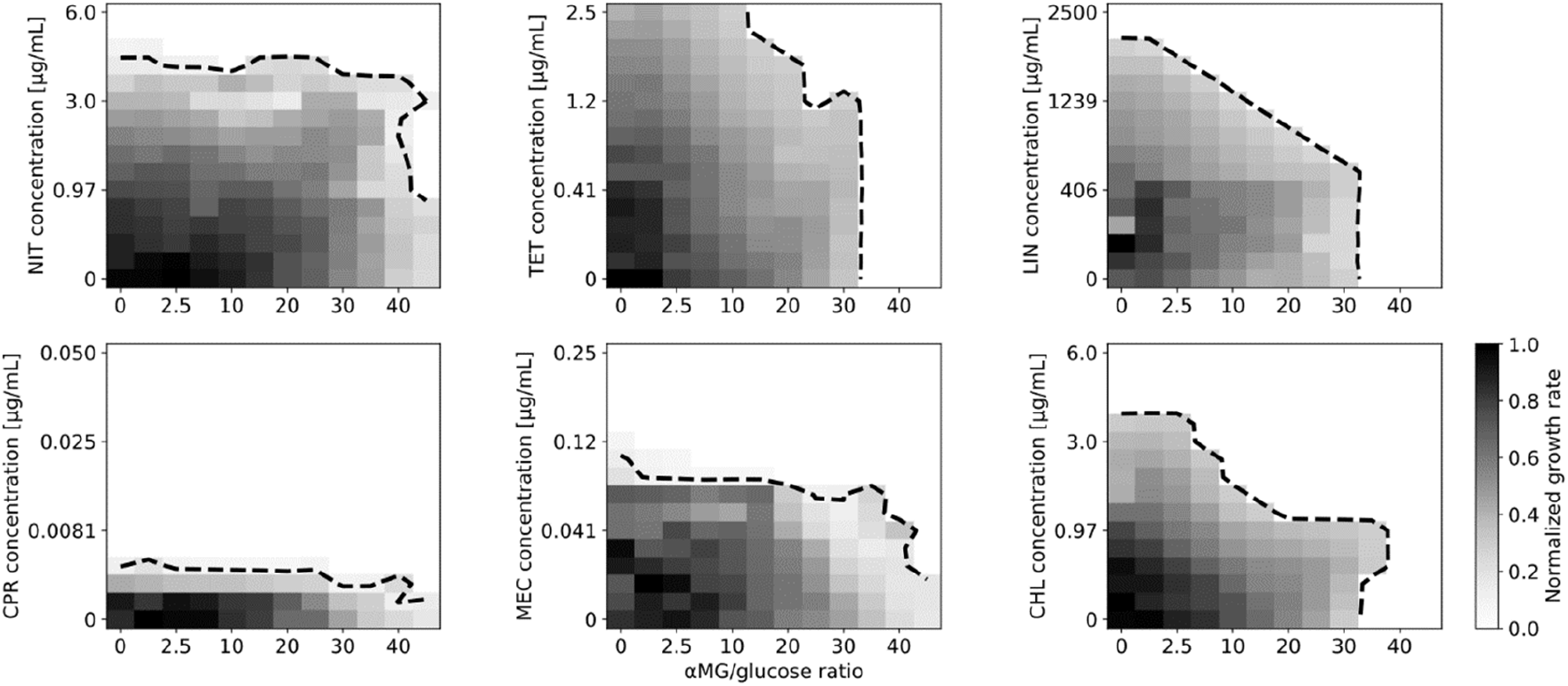
Effect of growth rate reduction by glucose limitation on susceptibility to diverse antibiotics. As Fig. 2B, for nitrofurantoin (NIT), tetracycline (TET), lincomycin (LIN), ciprofloxacin (CPR), mecillinam (MEC), and chloramphenicol (CHL). Lowering growth rate by glucose limitation via αMG does not lower susceptibility to these antibiotics as for TMP (cf. Fig. 2C).

**Supplementary Fig. 6.**
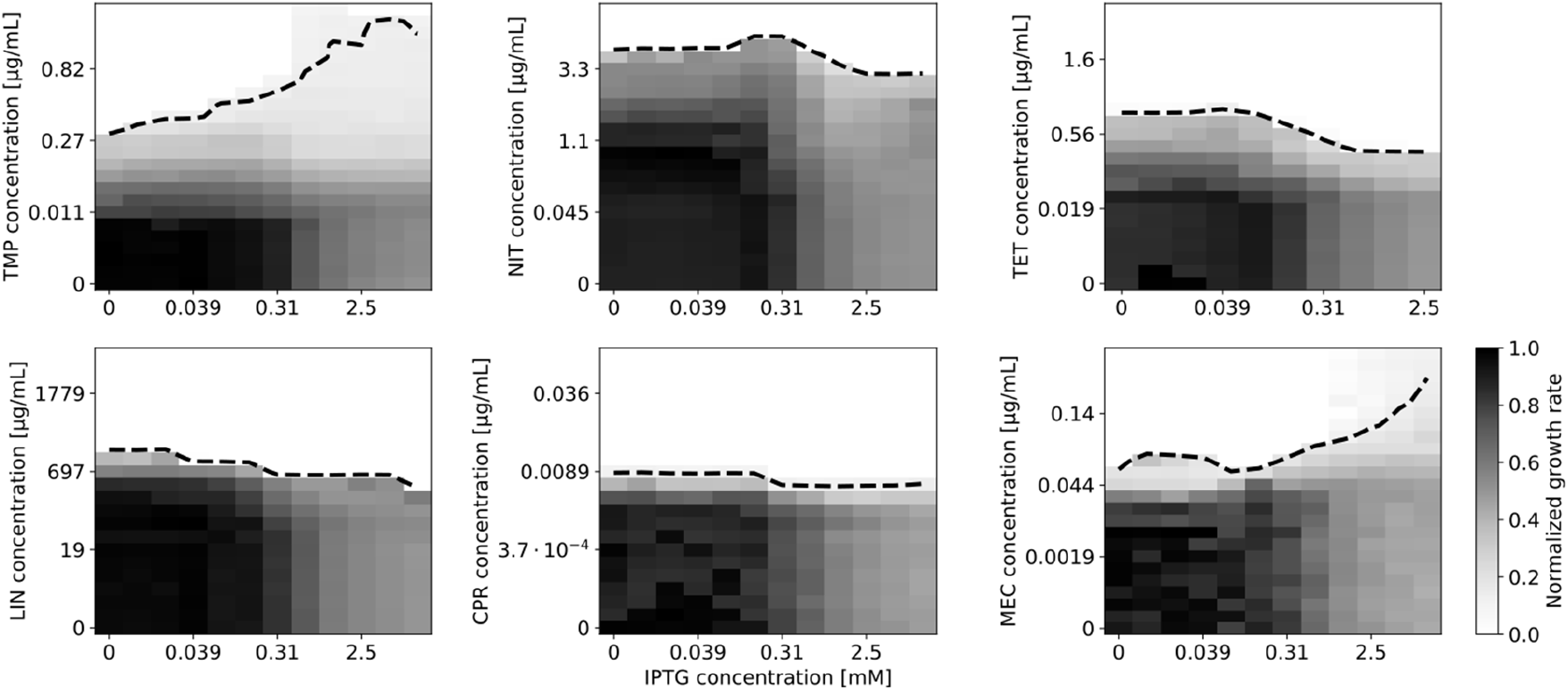
Effect of growth rate reduction by gratuitous protein overexpression on susceptibility to antibiotics. As Fig. 2E, for trimethoprim (TMP), nitrofurantoin (NIT), tetracycline (TET), lincomycin (LIN), ciprofloxacin (CPR), and mecillinam (MEC). Lowering growth rate by gratuitous protein overexpression lowers susceptibility to TMP and, to a lesser extent, to MEC (bottom right), but not for the other antibiotics (cf. Fig. 2F).

**Supplementary Fig. 7.**
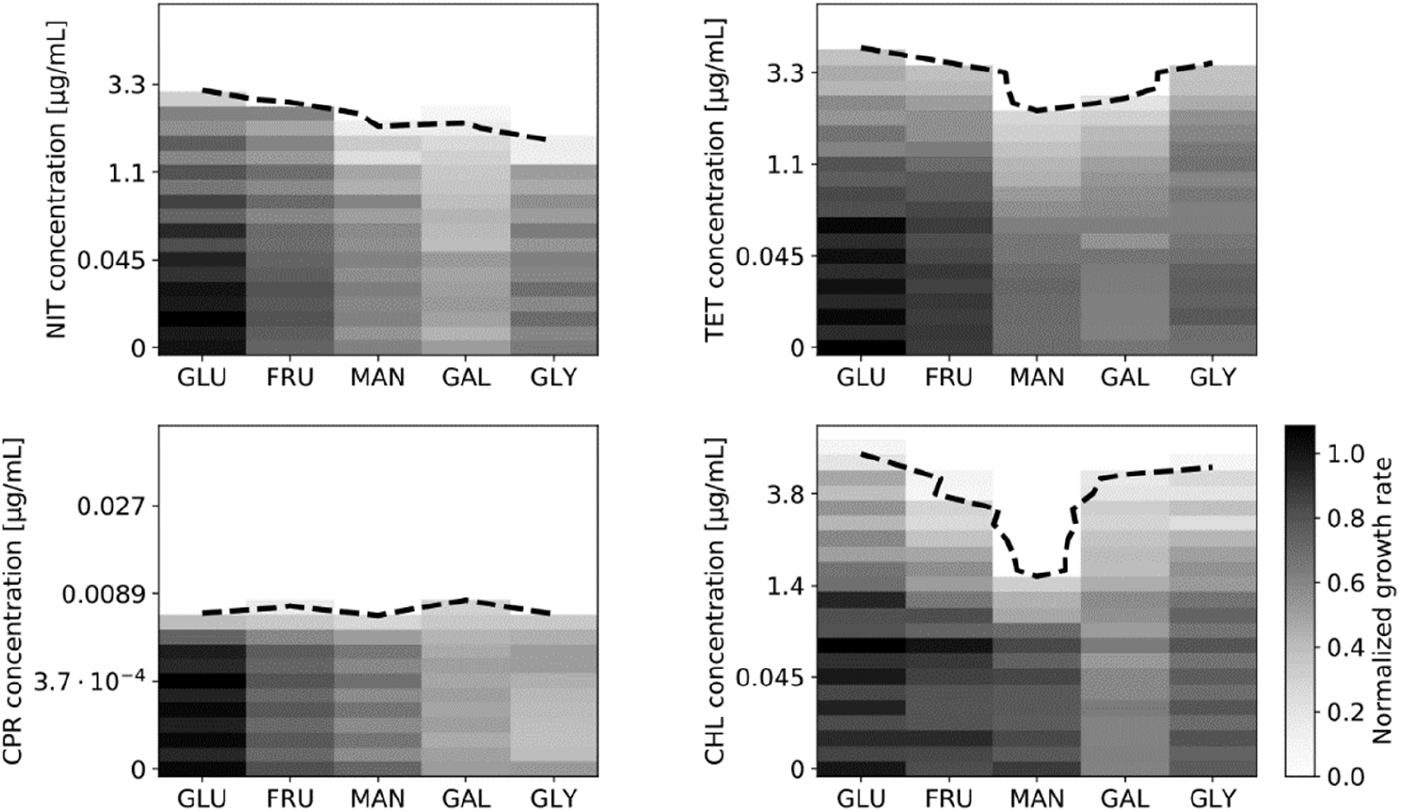
Effect of growth rate reduction by changing carbon source on susceptibility to antibiotics. As Fig. 2H, for nitrofurantoin (NIT), tetracycline (TET), ciprofloxacin (CPR), and chloramphenicol (CHL). Lowering growth rate via poorer carbon sources does not lower susceptibility to other antibiotics than TMP (cf. Fig. 2I). Data shown is the mean of three replicates. We also performed this assay for mecillinam (MEC), but excluded it from further analysis because – for unknown reasons – it consistently showed extremely noisy dose-response curves in this assay.

**Supplementary Fig. 8.**
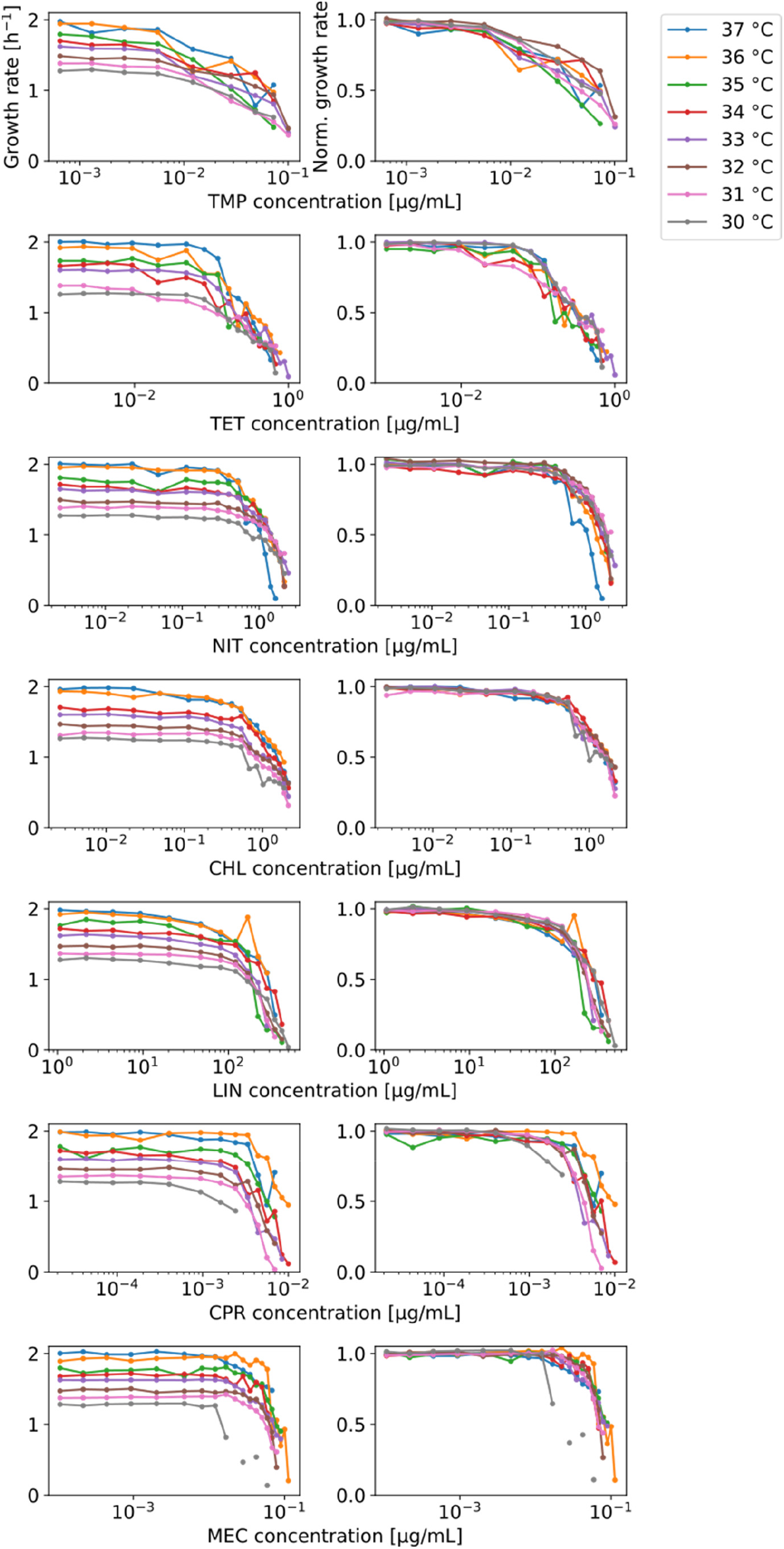
Lowering growth rate by changing temperature does not affect the shape of antibiotic dose-response curves. Left column: Growth rate versus drug concentration for eight different antibiotics at eight different temperatures as shown. Right column: Growth rate normalized to drug-free growth rate.

**Supplementary Fig. 9.**
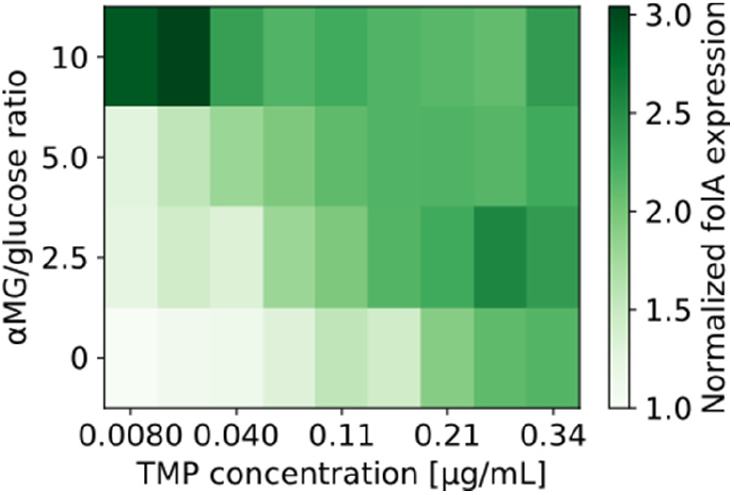
Expression level of folA as a function of trimethoprim and αMG/glucose ratio. Data from Fig. 2 in two-dimensional checkerboard plot.

**Supplementary Fig. 10.**
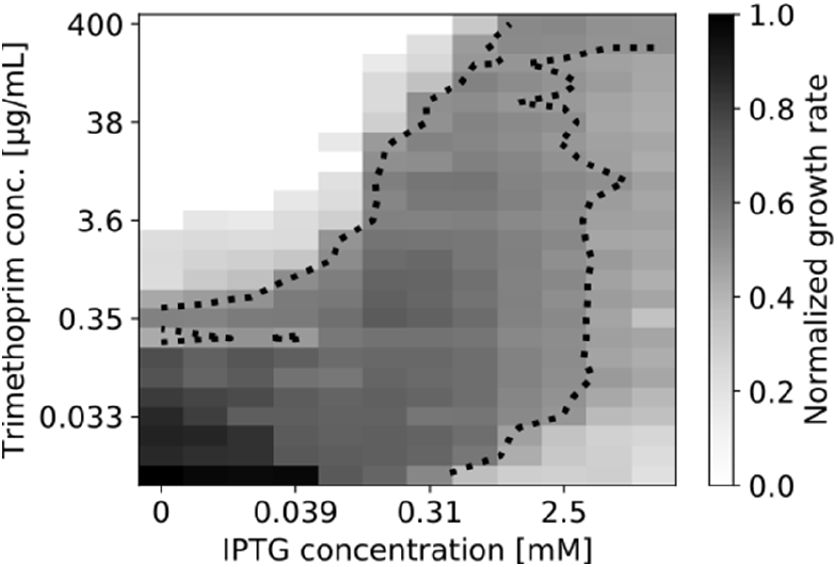
Growth rate decrease due to DHFR overexpression is rescued by trimethoprim. Normalized growth rate (gray scale) in a two-dimensional concentration gradient of IPTG and TMP. IPTG controls overexpression of folA (Methods). Dotted lines are contour lines at 50% growth inhibition. DHFR overexpression lowers growth rate but adding TMP at high IPTG concentrations partially rescues this phenotype: Growth rate increases with increasing TMP concentration. The increase in TMP IC_50_ resulting from DHFR overexpression confirms previous reports (Palmer & Kishony, 2014).

**Supplementary Fig. 11.**
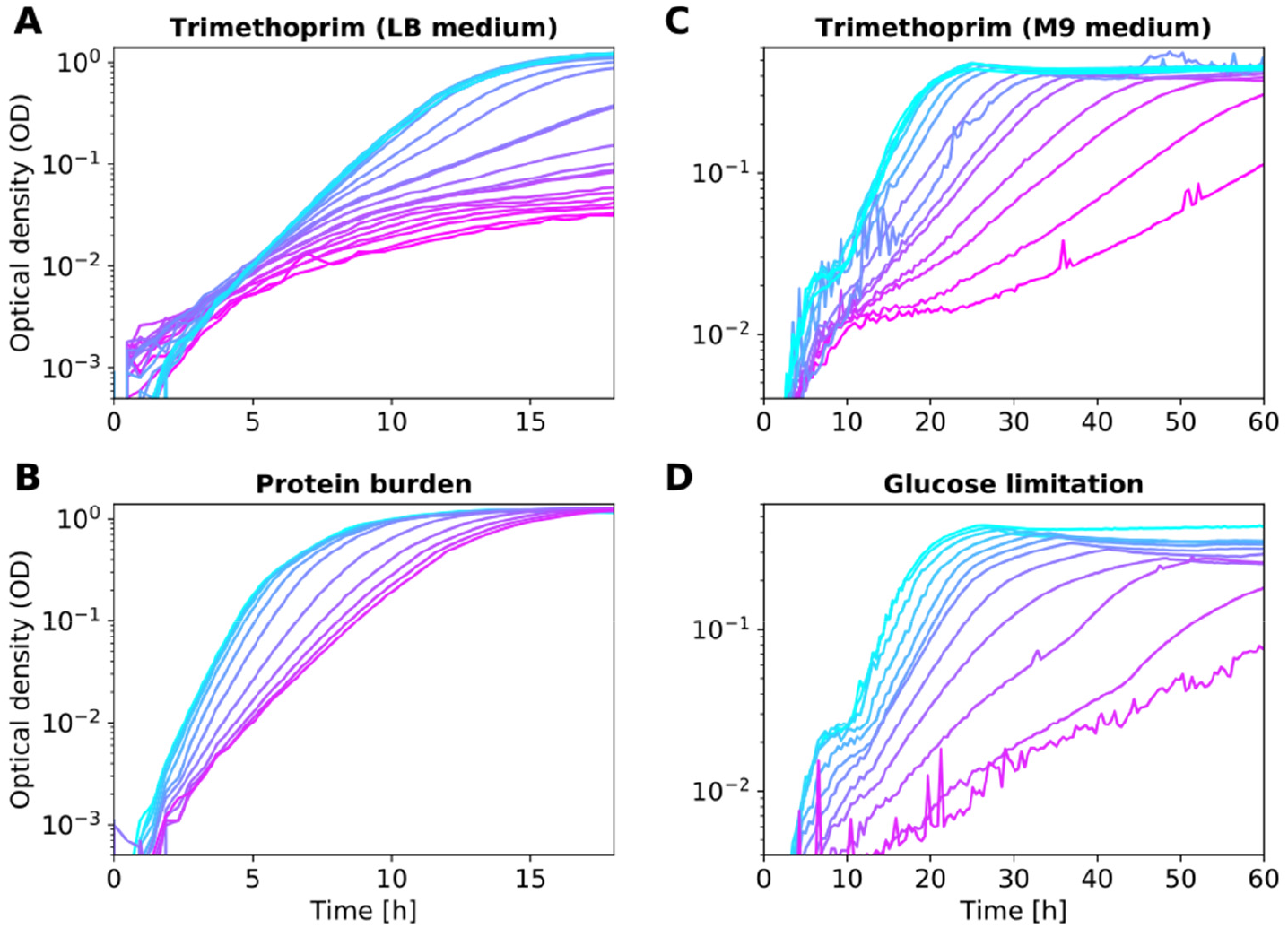
Bacterial growth curves from optical density measurements. Representative data for different ways of lowering the growth rate in different growth media. **(A)** TMP in LB medium. Growth curves underlying the data shown in the leftmost column of Fig. 2E; TMP concentration increases from cyan to magenta. **(B)** Gratuitous protein overexpression in LB medium; data correspond to Fig. 2D; IPTG concentration increases from cyan to magenta. **(C)** TMP in glucose (minimal M9) medium; data correspond to the leftmost column of Fig. 2B. **(D)** αMG in glucose (minimal M9) medium; data correspond to Fig. 2A; αMG/glucose ratio increases from cyan to magenta.

